# Bilingual Language Model for Protein Sequence and Structure

**DOI:** 10.1101/2023.07.23.550085

**Authors:** Michael Heinzinger, Konstantin Weissenow, Joaquin Gomez Sanchez, Adrian Henkel, Milot Mirdita, Martin Steinegger, Burkhard Rost

## Abstract

Adapting large language models (LLMs) to protein sequences spawned the development of powerful protein language models (pLMs). Concurrently, AlphaFold2 broke through in protein structure prediction. Now we can systematically and comprehensively explore the dual nature of proteins that act and exist as three-dimensional (3D) machines and evolve as linear strings of one-dimensional (1D) sequences. Here, we leverage pLMs to simultaneously model both modalities by combining 1D sequences with 3D structure in a single model. We encode protein structures as token sequences using the 3Di-alphabet introduced by the 3D-alignment method *Foldseek*. This new foundation pLM extracts the features and patterns of the resulting “structure-sequence” representation. Toward this end, we built a non-redundant dataset from AlphaFoldDB and fine-tuned an existing pLM (ProtT5) to translate between 3Di and amino acid sequences. As a proof-of-concept for our novel approach, dubbed Protein structure-sequence T5 (*<u>ProstT5</u>*), we showed improved performance for subsequent prediction tasks, and for “inverse folding”, namely the generation of novel protein sequences adopting a given structural scaffold (“fold”). Our work showcased the potential of pLMs to tap into the information-rich protein structure revolution fueled by AlphaFold2. *ProstT5* paves the way to develop new tools integrating the vast resource of 3D predictions, and opens new research avenues in the post-AlphaFold2 era. Our model is freely available for all at https://github.com/mheinzinger/ProstT5.

## Introduction

Large Language Models (LLMs) powered by Transformers^2^ have revolutionized Natural Language Processing (NLP) and have spawned ChatGPT^3,4^ affecting the daily life of millions. Adapting these techniques to protein sequences by equating words with amino acids and sentences with protein sequences started a wealth of new, powerful tools modeling proteins^5–13^. The success of these protein Language Models (pLMs) builds heavily on the rethinking of how to best leverage evolutionary information from large but unlabeled data. Instead of retrieving evolutionarily related proteins from large databases, pLMs extract meaningful features directly from protein sequences which are described by one-dimensional (1D) strings of 20 letters for the 20 common amino acids. The *knowledge* acquired by pLMs during pre-training is readily transferable to subsequent protein prediction tasks in the form of the hidden states of the pLMs (dubbed *embeddings*). For general-purpose pLMs, this so-called transfer learning succeeds for many aspects of protein prediction, including, for function prediction: gene ontology^14^, enzymatic function^15^, transport signals^16^, binding residues^17^, or subcellular location^18,19^, and for protein structure prediction: 2D^20^ and 3D structure^8^, fold classification^21,22^, or intrinsically disordered regions^23,24^. The same knowledge extracted by the pLM, can also be queried for protein design^25–27^, dynamic optimization^28–30^ or the inference of drug-target interactions^31^.

Concurrent with pLMs, *AlphaFold2*^32^ was able to predict highly-accurate protein structures to a degree near indistinguishable from experimental structure determination methods. By January 2024 accurate structure predictions (AlphaFold Protein Structure Database or AFDB^33^) are available for over 214 million protein sequences in UniProt^34^. This break-through opens up exciting possibilities to explore the dual nature of proteins - from their one-dimensional (1D) amino acid sequences to their unique three-dimensional (3D) structures.

Here, we propose leveraging pLMs to simultaneously model both modalities (1D and 3D). First, we encoded 3D structures as 1D strings (of tokens) to make them amenable to LLM techniques. Towards this end, we used the 3Di-alphabet introduced by the structure comparison method Foldseek^1^. Essentially, 3Di transliterates 3D coordinates into 1D strings of 20 letters, with one letter describing each residue’s (sequence position) interactions in a protein (the number is deliberately identical to that of natural amino acids (AAs)). This allows plugging-in highly optimized sequence search algorithms to compare 3D structures^35^. The same conversion allows feeding sequences of 3Di into a pLM. Besides learning to extract features from 3D structure, our solution invites switching between both modalities, i.e., translating from sequence to structure, and *vice versa* (Fig. 1). This opens new scientific avenues toward, e.g., inverse folding, structure-guided mutation effect prediction or remote homology detection.

**Fig. 1:**
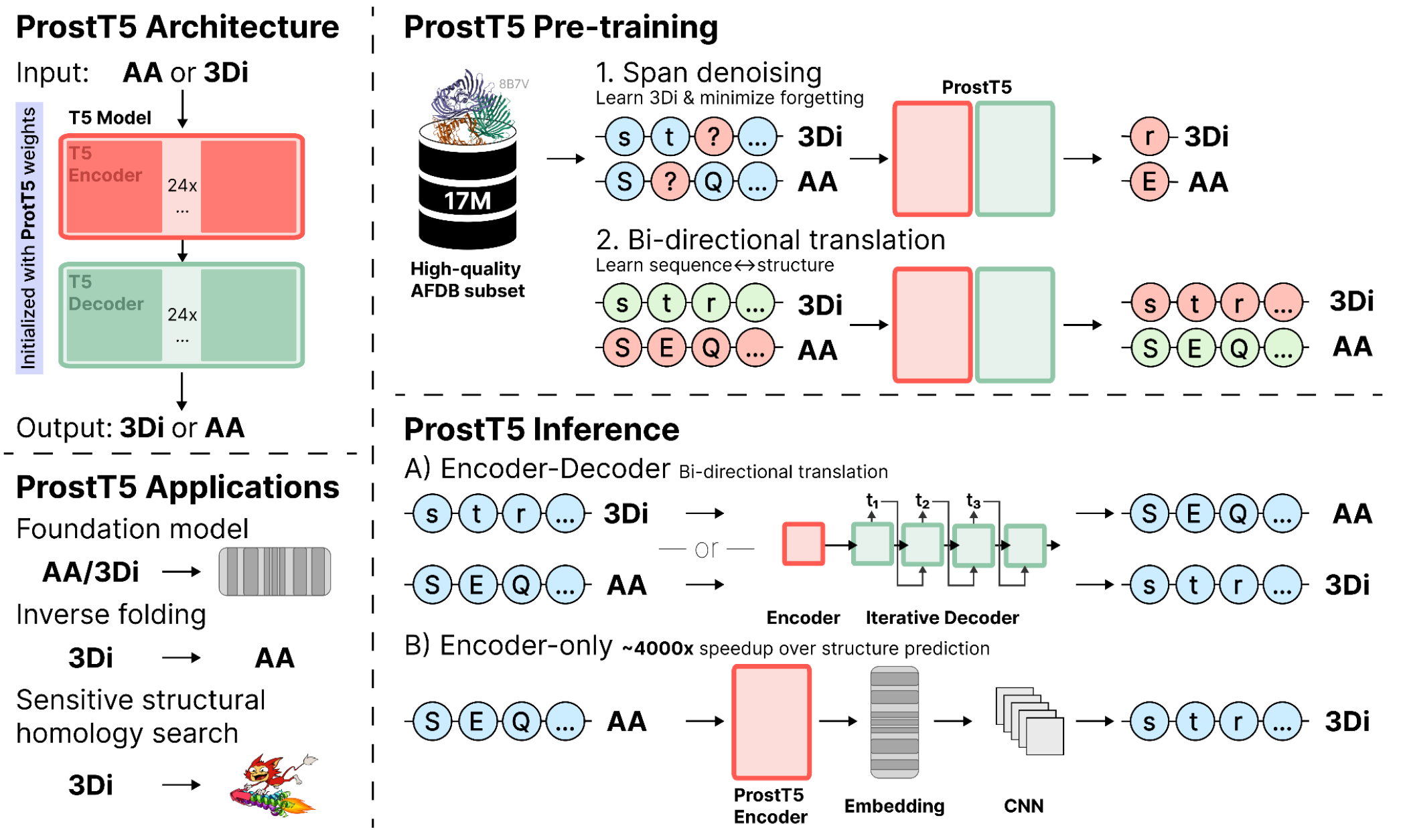
Sketch of ProstT5. *<u>Model architecture</u>:* ProstT5 is a T5-based encoder-decoder model initialized with weights of the ProtT5 model^6^. *<u>Pre-trainin</u>g:* Foldseek^1^ transferred protein 3D coordinates into 3Di tokens, i.e., 1D descriptions of 3D structure that assign each residue in a protein into one of twenty states described by a 1D string of letters. We used 17 million (17M) high-quality, non-redundant and diverse 3D predictions from AFDB^33^. ProtT5 was leveraged as an already pre-trained starting point for translating between 1D sequence (amino acids, AA) and 3D structure (3Di). Firstly, we applied the original pre-training objective of ProtT5 (span-based denoising) to both, AAs and 3Di, to teach the model the new 3Di tokens while avoiding catastrophic forgetting of AAs. Secondly, we continued to train the resulting model to translate between AAs and 3Di and *vice versa*. The final model, ProstT5 (Protein structure-sequence T5) extracts the information in its internal embeddings that can be input into downstream applications. This includes established feature extraction using only the encoder^6^, or bi-directional translation, either from AAs to 3Di (“folding”) or from 3Di to AAs (“inverse folding”). *<u>Inference</u>*: bi-directional translation from AA to 3Di (AA→3Di) or 3Di→AA can be conducted using either the encoder-decoder mode, necessitating token-wise decoder-inference or through an optimized inference mode, where 3Di tokens are directly predicted through a convolutional neural network from the encoder-embedding. The optimized 3Di inference mode results in a three orders of magnitude speedup over 3Di extraction from predicted protein structures (Fig. 2).

To quickly summarize our contributions, we:

- created a new dataset of diverse proteins with high-quality Alphafold2 3D structure predictions
- fine-tuned an existing pLM (ProtT5^6^) to translate between protein structure (3Di) and sequence. We dubbed the proposed model Protein structure-sequence T5 (ProstT5).
- demonstrated that ProstT5 can generate new protein sequences solely from their 3Di representation
- showed that 3Di sequences predicted by ProstT5 outperformed traditional sequence-based alignment methods in identifying distantly related proteins (*remote homology detection*) and offer structure-level search sensitivity to sequence-searches orders-of-magnitude faster compared to first predicting structures. We integrated the proposed 3Di prediction into the *Foldseek* webserver^1^.
- demonstrated that ProstT5 embeddings expand on its mono-lingual base-model ProtT5 in capturing aspects of protein structure.

## Results

### ProstT5 pre-training

We took the 3D structures for our ProstT5 data splits from a recently published resource^36^ which clusters *AlphaFold2*^32^ predictions in the AFDB^33^ using sequence- and structure similarity. After further filtering, e.g. for reliably predicted 3D structure (see Methods for more details), the remaining proteins were, on average, shorter than proteins in the PDB collecting experimental 3D structures (Fig. S1A; PDB^37^: 255 residues, compared to test: 206 and train: 238). The amino acid (AA) composition, however, was similar between reliable AlphaFold2-prediction and experiment (Fig. S1C). In stark contrast, the 3Di-tokens generated from the 3D structure (predicted or experimental) through *Foldseek*^1^ exhibited a severe class imbalance towards few over-represented tokens (Fig. S1D), in particular for the three states *v*, *d*, and *p* (we used lower- and upper-case letters for 3Di and AAs, respectively). These three tokens described over half the residues in our dataset. This imbalance was higher for our data than for the PDB at large. To better understand this finding, we mapped 3Di tokens from the PDB onto three secondary structure classes (Fig. S1B). This revealed a clear preference of some 3Di-tokens towards helix (v,l), strand (e,i,k,t,w,y) or other (d,p). In fact, 60% (12 out of the 20) of the 3Di-tokens had a clear preference (>70%) for one secondary structure class (helix, strand, or other).

Using the set *train17M*, we expanded ProtT5’s^6^ original pre-training objective (span-based denoising^38^ of AAs) to cover both, AA- and 3Di-sequences, which yielded 2*17M=34M samples. When the validation loss plateaued, the actual training on the translation from AAs to 3Di and *vice versa* commenced. Before clear convergence (loss still decreasing slightly), we stopped training due to the decreasing tradeoff between cost (energy, i.e., computing resources) and improvement.

### 3600-fold faster structural remote homology detection at same sensitivity

Inputting amino acid sequence, ProstT5 can generate (predict) 3Di states for each residue, and these in turn can be used as input to alignment methods. Thereby, ProstT5 readily turns into a tool for remote homology detection. For comparison, we replicated the *Foldseek*^1^ benchmark. We replaced structure-sequences (3Di) derived from experimental structures in the Foldseek benchmark (40% non-redundant version of SCOPe^39^) by 3Di strings predicted by ProstT5 from corresponding amino acid sequences. We compared the sensitivity up to the first false-positive (FP) on the levels of *family*, *superfamily* and *fold* for Foldseek applied on 3Di-sequences derived from experimental data (Fig. 2: Foldseek(3Di)) against the ProstT5-generated 3Di strings (Fig. 2: Foldseek(p3Di)). We compared traditional sequence alignments through *MMseqs2*^35^. Using predicted 3Di states, ProstT5 (ROC-AUC on the level of super-family: 0.45) reached performance levels close to experimental structures (ROC-AUC=0.49) and vastly outperformed traditional sequence-based searches (ROC-AUC=0.06). On top, Foldseek using 3Di strings from experiments needs C-alpha coordinate information to improve sensitivity. When disabling this refinement, sensitivity dropped below the value reached by ProstT5 predictions, and when adding C-alpha coordinate information to predictions, it increased slightly over the experimental value (data not shown).

**Fig. 2:**
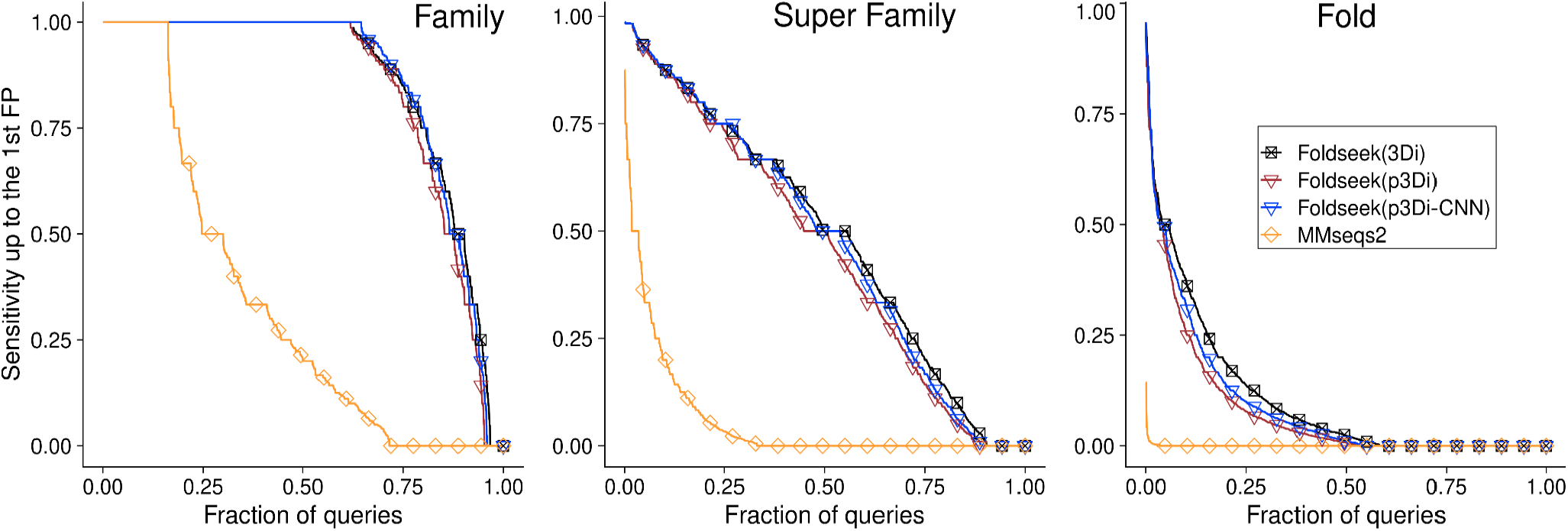
Successful remote homology detection with predicted 3Di. We replicated the Foldseek benchmark^1^ on SCOPe40^39^ using 3Di strings generated either by ProstT5 (Foldseek(p3Di)) or a CNN trained on top of ProstT5’s encoder (Foldseek(p3Di-CNN)) and compared the sensitivity up to the first false positive (protein with different classification) with the performance of Foldseek on experimental structures (Foldseek(3Di)). For all three levels (from fine-grained family level on the left, over the superfamily level, to the coarse-grained level of *fold*), ProstT5-predicted 3Di strings sufficed to almost reach the performance of PDB structures while significantly outperforming traditional sequence alignment (*MMseqs2*^35^).

To provide a more lightweight alternative for generating 3Di, we trained a two-layer CNN to predict 3Di states directly from amino acid embeddings derived only from the encoder of ProstT5 (Fig. 2: Foldseek(p3Di-CNN)). This avoided unnecessary inference slow-down from the decoder’s auto-regressive generation at high remote homology detection sensitivity (ROC-AUC=0.47 on the super-family level). We also compared the 3Di classification accuracy of identical CNNs trained on different embedding sources, i.e., ESM-2, ProtT5, Ankh, and ProstT5, which highlighted the benefit of ProstT5’s expanded structure-sequence pre-training compared to only AA pre-training (SOM Fig. S3). Similar to AlphaFold’2s predicted reliability pLDDT, users can apply a threshold on the CNN output to focus either on high precision (high threshold) or on high coverage/recall (low threshold, Fig. S5).

This enables sensitive whole-proteome annotation within minutes without requiring time- and resource-consuming structure prediction. For instance, predicting 3Di tokens for the proteome of *Methanocaldococcus jannaschii* took 40min 6s (±4min 38s) on CPU (Macbook M1 2020) and 43.5s (±2s) on GPU (Nvidia RTX A5000), compared to the 48h structure prediction time on GPU (or to an extrapolated 112 day prediction time on the same Macbook CPU) for the proteome with an optimized ColabFold workflow^40^. Utilizing predicted 3Di tokens through ProstT5+Foldseek for remote homology detection thus enables an over three orders of magnitude speedup over ColabFold-based generation of 3Di strings. Our unoptimized prediction workflow paves the way for sensitive protein annotation not only for proteome-wide scale, but even for metagenomic scale datasets.

### Bilinguality improved structure encoding

As before^6^, we benchmarked the resulting pLM ProstT5 on representative prediction tasks. We began with secondary structure prediction as arguably the best understood proxy problem. As usual for pLMs, we extracted the hidden states of the encoder’s last layer, and input these vectors, dubbed *<u>embeddings</u>*, into the subsequent, 2nd-step supervised prediction tasks. Given ProstT5’s bilinguality, we could derive embeddings for both AA- and 3Di-sequences. The embeddings were input into a convolutional neural network (CNN) to classify each residue into helix, strand, or other (Fig. 3A). We used *biotrainer*^41^ together with FLIP^42^ to replicate the training setup of previous work^6^, i.e., we optimized a two-layer CNN on the *NetSurfP-2.0* training data^46^ and measured performance on three test sets (*CASP12*^43^, *CASP14*^44^ and *NEW364*^6^). The difference in performance (measured by the three-state per-residue accuracy, Q3^47^) between the three sets estimated the error (lower limit: CASP14, upper limit: NEW364; dot between: CASP12). The existing state-of-the-art (SOTA) general–purpose sequence-based pLMs (ProtT5, ESM-2^8^ and Ankh^12^) appeared to perform alike with ProtT5 being slightly worse. Even without leveraging 3Di-information (ProstT5(AA) used only AA input), bilingual ProstT5 improved over its base model ProtT5 (Fig. 3A). When inputting 3Di-sequences derived from experimental structures (ProstT5(3Di)), Q3 approached 90% on *NEW364*. On the one hand, this came close to the upper bound given by the agreement between different experimental determinations for the same protein^48^. On the other hand, this comparison was crucially circular: inputting 3D structure to predict secondary structure is meaningless, in practice. This is exemplified by reaching performance competitive to SOTA even when only using one-hot-encoded experimental 3Di-sequences (OHE(3Di)). In contrast, when using 3Di-sequences predicted by ProstT5 (labeled *ProstT5(p3Di)*), Q3 dropped below the base model ProtT5. When inputting concatenations of sequence and structure embeddings performance increased numerically, but not statistically significantly (Fig. S2A).

**Fig. 3:**
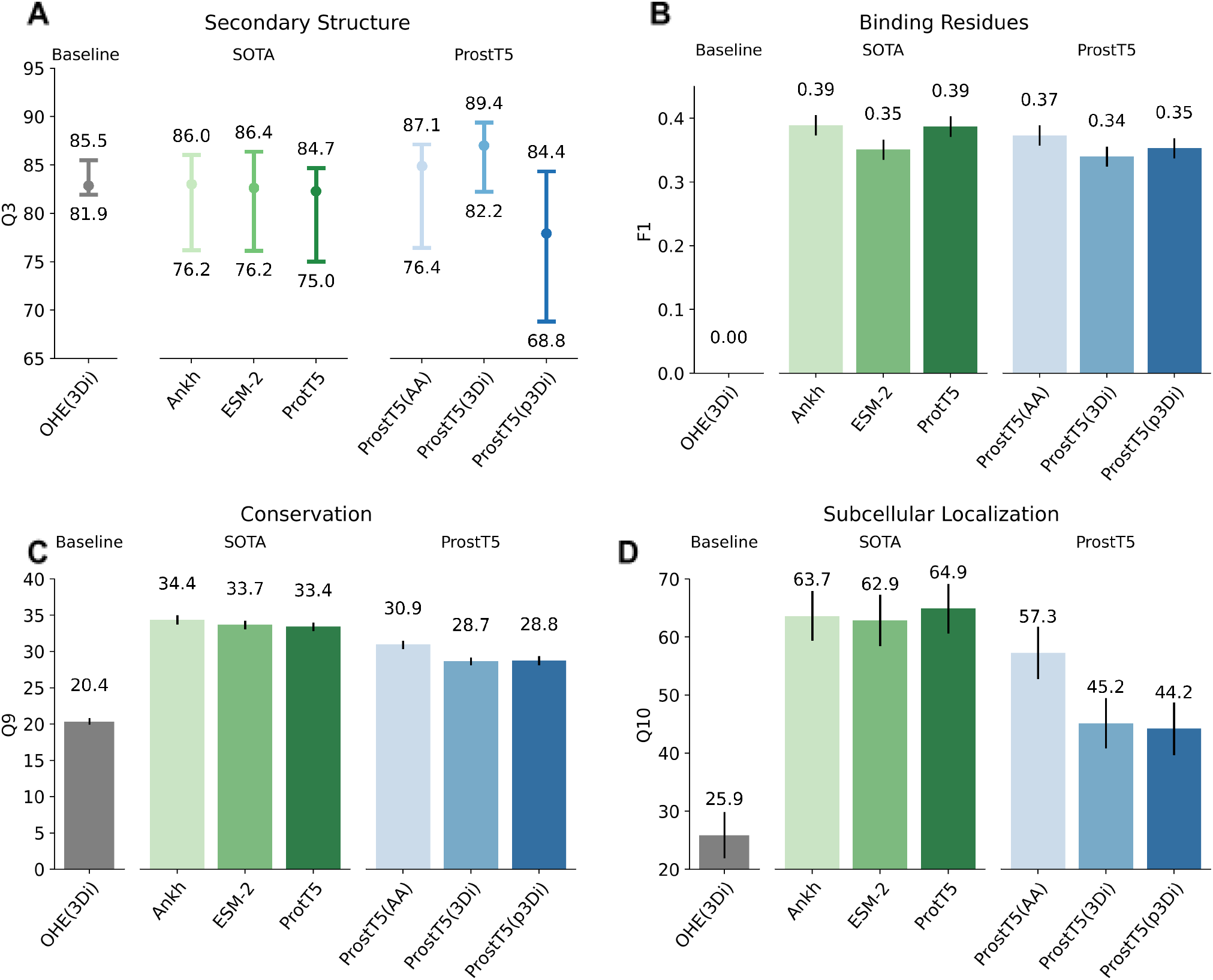
Protein prediction tasks exclusively using pLM embeddings. We probed the relevance of the information learned by ProstT5 by inputting its embeddings into subsequent supervised prediction methods, as introduced before^5^. In particular, we compared ProstT5 to SOTA general purpose pLMs using only amino acid sequences (ProtT5^6^, Ankh^12^, and ESM-2 (3B)^8^) on four different prediction tasks, namely the per-residue prediction of secondary structure (<u>A</u>: performance: Q3, three-state per-residue accuracy; data sets: middle: *CASP12*^43^, lower bar: *CASP14*^44^, upper bar: *NEW364*^6^; note: since each set is supposed to measure performance, the difference between these provided an error estimate), binding residues (<u>B</u>: performance: F1; data: *testSet300*^17^), conservation (<u>C</u>: performance: Q9, nine-state per-residue accuracy; data: ^45^), and the per-protein prediction of subcellular location (<u>D</u>: performance: Q10, ten-state per-protein accuracy; data: *setHARD*^18^). As a baseline, we also probed the information content readily available from one-hot-encoded 3Di-sequences (OHE(3Di)). For panels B-D, the bars mark the 95% confidence interval, i.e., ±1.96 * standard errors, estimated via bootstrapping.

### ProstT5 is not a general-purpose pLM

We tried to avoid *catastrophic forgetting*^49^ of what ProtT5 had extracted from pre-training on protein sequences during fine-tuning by continuing de-noising on amino acid sequences and bi-lingual translation. Nevertheless, some information might have been lost during this process as shown by a clear decrease in predicting subcellular location when inputting exclusively amino acid sequences (Fig 3C: ProtT5 vs ProstT5(AA)). Other tasks, such as the prediction of binding residues (Fig 3B) or conservation (Fig 3D), showed a similar trend, albeit with lower performance drop. One-hot-encoding of 3Di (OHE(3Di)) also appeared less useful for those tasks. Concatenating amino acid embeddings from ProtT5 and ProstT5, can compensate for this, and can even lead to a numerical improvement as shown for binding residue prediction (Fig. S2B).

### CATH classification of proteins

We also assessed performance beyond single residues by benchmarking ProstT5-embeddings on the classification of proteins into structural classes (often referred as *folds*) using embedding-based annotation transfer (EAT^21^; replacing sequence-similarity by the Euclidean distance in embedding space to transfer annotations from a lookup database to a query protein). To simplify comparability, we replicated existing benchmarks^21^ on CATH^50^ which uses structural similarity to capture evolutionary and functional relationships of proteins beyond sequence similarity. Given that ProstT5 can generate embeddings from either sequence (ProstT5(AA)) or structure input (or both), we benchmarked both different input modes (predicted 3Di: ProstT5(p3Di), experimental 3Di: ProstT5(3Di)). All improved over ProtT5, ESM-1b and Ankh (Table 1). Compared to ProtT5, embeddings from amino acids (ProstT5(AA)) mostly improved for the CATH levels of architecture (CATH-A, Table 1) and topology (CATH-T, Table 1), while embeddings from 3Di states (ProstT5(3Di) and ProstT5(p3Di)) improved most for the fine-grained classification of homologous superfamilies (CATH-H). This orthogonal information can be leveraged by concatenating the embeddings from amino acid- and predicted 3Di-sequences (ProstT5(cat)) improving over either method at all CATH levels. We also benchmarked the effect of optimizing ProstT5-embeddings using contrastive learning^21^ and while overall performance improved through task-specific optimization, the general trends remained the same (SOM: Table S1).

**Table 1:** Classification of proteins into CATH hierarchy (folds)*

|  | Method/Input | CATH-C | CATH-A | CATH-T | CATH-H | Mean |
| --- | --- | --- | --- | --- | --- | --- |
| Baseline | Random<sup>‡</sup> | 29±6 | 9±4 | 1±2 | 0±0 | 10±3 |
| HBI | MMSeqs2<sup>‡</sup> | 52±7 | 36±6 | 29±6 | 35±6 | 38±6 |
| EAT<br>unsupervised | ESM-1b<sup>‡</sup> | 79±5 | 61±6 | 50±7 | 57±8 | 62±7 |
|  | Ankh | 84±5 | 69±6 | 60±7 | 67±8 | 70±6 |
|  | ProtT5<sup>‡</sup> | 84±5 | 67±6 | 57±6 | 64±8 | 68±6 |
|  | ProstT5(AA) | 85±5 | 74±6 | 64±6 | 69±7 | 73±6 |
|  | ProstT5(p3Di) | 85±5 | 71±6 | 60±7 | 73±7 | 72±6 |
|  | ProstT5(3Di) | **90±4** | **77±6** | **65±6** | **75±7** | **77±6** |
|  | ProstT5(cat) | 88±4 | 74±6 | 65±7 | 74±7 | 75±6 |

\* **Performance:** accuracy for predicting CATH<sup>50</sup> levels (from coarse- to fine-grained: C, A, T, H) by transferring annotations from a lookup set to a strictly non-redundant set of queries taken from<sup>21</sup>. The column *Mean* marked the average over the four performance values. Values of methods marked by the symbol <sup>‡</sup> were taken from the literature<sup>21</sup>. **Methods:** *Baseline:* Random transfer of labels by randomly picking a protein; *HBI (homology-based inference):* MMSeqs2<sup>35</sup> used single sequence search to transfer the annotation of the hit with the lowest *E*-value; *EAT-unsupervised:* embedding-based annotation transfer using the shortest Euclidean distance measured in embedding space of unsupervised pLMs ESM-1b<sup>7</sup>, ProtT5 and ProstT5 with different inputs (AA=amino acid sequence input, p3Di=3Di predicted by ProstT5, 3Di=3Di from experimental structures, cat=concatenation of AA and p3Di). Error bars mark the 95% confidence interval with $\pm 1.96$ standard errors. Bold numbers mark the highest numerical values. Note that the error bars were so high that statistically significant differences were only observed between *random* and *HBI*, as well as those between *EAT* and everything else (although the lowest end of pLMs and ProstT5(cat), just differed significantly within the 95% confidence interval).

### *Inverse Folding*: creating new diverse protein sequences with similar structure

The *bilingual* nature of ProstT5 (AA→3Di and 3Di→AA) suggested creating diverse sets of never-before seen amino acid sequences that adopt a particular structure (as described by its 3Di-tokens). As pairs of proteins with diverged sequences may adopt similar 3D structures^51,52^, we measured success in creating new sequences through the similarity in predicted 3D structure between the *groundtruth* (3D predicted by AFDB^33^) and *predictions* (3D structure predicted by ESMFold^8^ for ProstT5-generated sequences). As our model assigns probabilities to a sequence of amino acids given some conditioning upon context (3Di), we used the validation set to compare the influence of hyper-parameters (incl. beam-search^53^, nucleus sampling^54^, and top-k sampling^55^) on the translation (3Di→AA) and its quality by modulating the probability assigned to a particular sequence during sequential decoding (SOM: Table S3). For the final test set evaluation, we chose a configuration providing a trade-off between similarity (to the native) in terms of structure and unsurprising sequence (proxied by Kullback-Leibler divergence between the amino acid distribution in UniProt and the sequences generated^56^). We measured structural similarity (ESMFold prediction of generated sequences vs. AlphaFold2 prediction of the native sequence) by three scores: lDDT^57^,TM-score^58^ and RMSD (as implemented in ^58^).

First, we established an upper bound for performance by comparing ESMFold to AlphaFold2 predictions for the native sequence (Table 2: *Native/ESMFold*). Although ProstT5 generated sequences with, on average, as little as 21% PIDE (percentage pairwise sequence identity) to the native protein, these were all predicted to adopt similar structures (average lDDT=72), the *de facto* standard for inverse folding, i.e., graph-based *ProteinMPNN*^59^, succeeded in generating sequences close to this upper-bound (lDDT(ProteinMPNN)=77 vs. lDDT(Native/ESMFold)=78). However, the amino acid distribution of ProstT5-generated sequences was closer to the native distribution as measured by entropy (Table 2).

**Table 2:** Inverse folding comparison*.

|  | *Native/ESMFold* | *ProteinMPNN* | *ProstT5* | *ProstT5(RoundTrip70)* |
| --- | --- | --- | --- | --- |
| *IDDT* ↑ | 0.78±0.01 | **0.77±0.01** | 0.72±0.01 | 0.73±0.01 |
| *RMSD* ↓ | 2.55±0.01 | **2.61±0.01** | 2.90±0.01 | 2.81±0.01 |
| *TM-score* ↑ | 0.62±0.02 | **0.61±0.02** | 0.58±0.02 | 0.60±0.02 |
| *PIDENT* | 100±0 | **29.6±1** | 21.9±0.9 | 22.4±0.9 |
| *Entropy* ↓ | 0.13±0.01 | 0.39 +/- 0.03 | 0.20±0.01 | **0.19±0.01** |

\* Performance: structural similarity of ESMFold<sup>8</sup> and AlphaFold2<sup>32</sup> predictions for native (Natural/ESMFold) and generated sequences in our test set. Sequences were generated using ProteinMPNN, ProstT5 and a filtered version of ProstT5 (ProstT5(rTrip70)) which uses the intrinsic back-translation of the model to filter by sequence similarity between native 3Di sequences and their counterpart predicted from generated AA sequences (3Di→AA→3Di). We generated AA sequences either until convergence (defined as ≥70 percentage pairwise sequence identity - PIDE - for 3Di letters) or after maximally ten attempts (to conserve resources). Single-sequence based ESMFold predictions for generated sequences were compared against the native ground-truth predicted by AlphaFold2 using IDDT<sup>57</sup>, RMSD, TM-score<sup>58</sup>, PIDE, and entropy (KL-divergence between the AA distribution in UniProt and the generated sequences). Error bars indicate 95% confidence intervals estimated from 1000 bootstrap samples. Arrows next to metrics indicate whether higher (↑) or lower (↓) values are better. For PIDENT applied to inverse folding, it is not clear whether higher is necessarily better.

Motivated by the success of ProstT5-predicted 3Di-sequences for remote homology detection (Fig. 2), we also probed whether ProstT5-generated backtranslations (3Di→AA→3Di), provided any indication for the quality of the inverse folding (new sequences for given structure). Towards this end, we correlated the structural similarity (lDDT) of native sequences predicted by AlphaFold2 and ProstT5-generated sequences predicted by ESMFold against PIDE between the native 3Di-sequence and the 3Di-sequences generated from the translated AA sequence (Fig. S6A). We observed a high correlation (Spearman’s R of 0.64) for what we referred to as the *roundtrip accuracy*, i.e., translating from native 3Di to AAs which were then translated back into 3Di for comparison with the starting point (native 3Di). We further applied this idea to constrain generated sequences, i.e., we generated AA sequences until reaching a *roundtrip accuracy* ≥70% and retained only the candidate with the maximal *roundtrip accuracy*. Even when limiting this to ten for saving resources, we already observed a minor, yet consistent improvement for all metrics (Table 2 - ProstT5(rTrip70), Fig. S6B). Sequences generated by *ProstT5* and *ProteinMPNN* agreed well in their predictions (Fig. S6C, Spearman’s R of 0.52). For the proteins in our test set, there was no difference in the inverse folding performance between those from clusters and those that remained singletons (Fig. S6). We cherry-picked cases for which both models (ProstT5 and ProteinMPNN) generated sequences resulting in high-quality structures (Fig. 4A-B) but we also investigated cases for which either model failed (Fig. 4C and Fig. 4D, ProstT5>ProteinMPNN and ProstT5<ProteinMPNN, respectively).

**Fig. 4:**
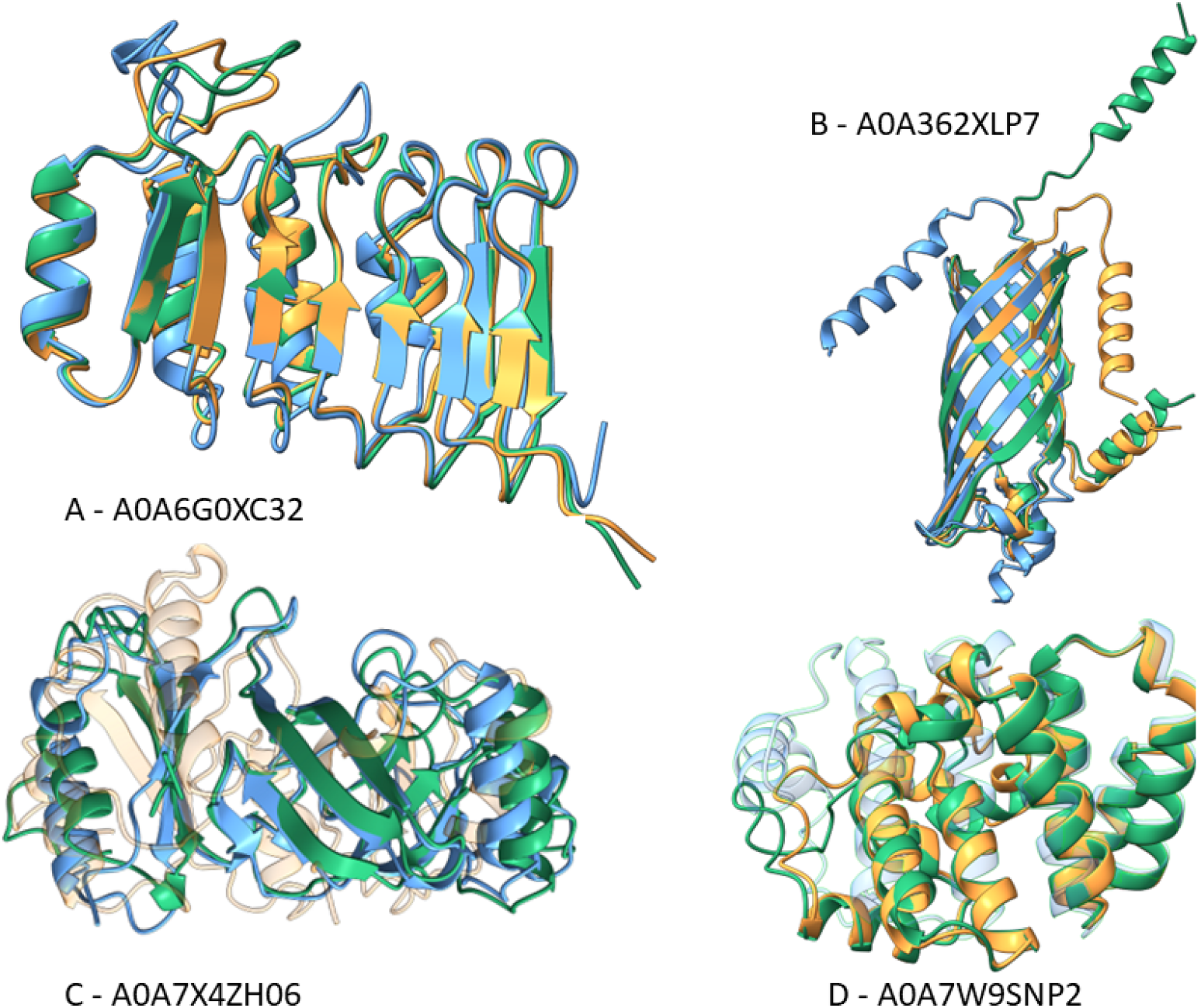
Inverse folding examples. We manually picked four examples from our test set for which both (A and B), ProstT5 and ProteinMPNN, only ProstT5 (C) or only ProteinMPNN (D) generated sequence with high structure similarity to their natural counterparts. Structures colored in green show the AlphaFold2 predictions (considered ground-truth), blue and orange depict ESMFold predictions of ProstT5- (blue) and ProteinMPNN-generated (orange) sequences, respectively. We picked examples such that they show diversity in their structural composition (beta-strands and alpha-helices) and their place of action (transmembrane (B) vs soluble (A,C,D)). Both methods can produce proteins that share only little sequence but high structural similarity to their native counterparts (A & B: lDDT of 76-95 or RMSD of 1.1-2.6 at 23-44% sequence similarity) but there are also cases where only one of them succeeds (C: ProstT5(lDDT)=68 vs ProteinMPNN(lDDT)=34; D: ProstT5(lDDT)=56 vs ProteinMPNN(lDDT)=75). For better visibility, we increased transparency for cases with poor structural superposition (C: ProteinMPNN, D: ProstT5).

### Speed

The time required to generate embeddings from the ProstT5 encoder, e.g., to predict 3Di states directly via a CNN, is identical for ProtT5 and ProstT5 due to the analogous network architecture^6^. Generating embeddings for the human proteome from both the ProstT5 and the ProtT5 encoder requires around 35m (minutes) or 0.1s (seconds) per protein using batch-processing and half-precision (fp16) on a single RTX A6000 GPU with 48 GB vRAM. The encoder-decoder translation is comparatively slow (0.6-2.5s/protein at an average length of 135 and 406, respectively) due to the sequential nature of the decoding process which needs to generate left-to-right, token-by-token. We only used batch-processing with half-precision and left further speed-ups via LLM-specific optimizations^60^ to future work.

## Discussion

### Standing on the shoulders of giants

The avalanche of recent progress in NLP was triggered by the introduction of attention^61^ paired with excellently scaling Transformers^2^. Mining this breakthrough requires models with billions of free parameters trained on gigantic data sets. In computational biology Transformers have been largely limited to training large language models (LLMs) on amino (AA) and nucleic acid sequences^62,62^. *AlphaFold2*^32^ with its over 200 million predictions (AFDB^33^) has changed this limitation to sequence-only, non-structure data fundamentally.

By enabling scalable structure search at the speed of sequence comparison, *Foldseek*^1^ opened the door to leveraging structure predictions. Foldseek needed to harvest several solutions to make this possible, the arguably most important was to map the 3D coordinates for each residue in a protein structure to one of the 20 so-called 3Di tokens through a vector-quantized variational autoencoder (VQ-VAE^63^). The resulting conversion from structure (3D) to sequence (1D) allows Foldseek to leverage highly optimized sequence comparison tools^35^ to compare 3D structures. Given the impressive success of Foldseek, we postulated that 3Di sequences contain enough information to train an LLM with the objective to translate from structure to sequence.

### Sampling from proteins’ Janus face

Janus is the double-faced Roman god of duality and transitions. Here, we combined two major ‘“*faces*” of proteins, namely sequence and structure (obtained from AlphaFold2 predictions, i.e., AFDB→3Di) by fine-tuning the sequence-based protein Language Model (pLM) ProtT5^6^ on 17M proteins, each with L (protein length) doublets of 3Di- and AA-sequences. Similar to the combination of images or movies with text^64^, we merged protein sequences (AA) and structures (3Di) into a single model, a new pLM, dubbed <u>ProstT5</u>. This increased the flexibility for querying knowledge stored in the weights of the pLM through expertise going beyond embedding extraction. Despite notable exceptions^9,25^, established pLMs are mostly limited to feature extraction from encoder-style Transformers^6–8^. Instead, T5’s encoder-decoder structure allows ProstT5 to additionally act as a conditional Language Model that assigns probabilities to a sequence of tokens given some conditioning context. We conditioned upon structure (3Di) to generate sequences (AAs) and *vice versa*. This bilingual translation capability opens many possibilities. For instance, ProstT5 enables direct 3Di predictions from AA sequences. In fact, the 3Di predictions were so accurate that when input into Foldseek, they allowed the identification of structural similarity between extremely divergent proteins (*remote homology detection*) almost at the level of experimental structures (Fig. 2). ProstT5 reached this impressively high level directly, without having to predict structure which greatly reduced time- and compute-resource requirements.

This paves the way for high-throughput remote homology detection at structure comparison sensitivity. Given that the sampling capacity of the ProstT5 decoder, i.e., length-variability or distribution over predictions, is not needed, we replaced it by a two-layer CNN trained to predict 3Di states directly from the encoder’s amino acid embeddings. This avoids unnecessary inference slow-down from the decoder’s auto-regressive generation while maintaining high remote homology detection sensitivity (Fig. 2). With this modification, we can predict 3Di sequences on average for ten proteins per second for the human proteome on a single GPU. Despite reaching only levels of classification accuracy of 41-65% (Fig. S3; Q20), predicted 3Di sequences still excelled at remote homology detection, presumably because mistakes mostly confused 3Di states coding for similar (secondary) structure motifs (Fig. S1B) that are likely to substitute each other (Fig. S4A vs Fig S4B).

Inverting the translation task (from 3Di→AA), ProstT5 successfully accomplished *inverse folding*, i.e., *creating* unknown proteins with likely similar structures and different sequences (Table 2, Fig. 4, Fig. S6). Although not reaching the *de facto* SOTA toward this end, namely ProteinMPNN^59^, which uses a graph-based, message-passing network for this task, our proof-of-concept already reached an average lDDT of 72 and even outperformed ProteinMPNN for some cases (Fig. 4C, S6C). This, at least, suggested some complementary of the two approaches. On top, our solution seemed to create more diversity.

More interesting applications emerge when combining both translation directions in series. We showcased the usefulness of stringing together both directions by using ProstT5 to assess the quality of its own predictions generated during inverse folding (Fig. S6B). First ProstT5 generated novel AA sequences conditioned upon adopting any desired structural template (here given by the 3Di sequence from AlphaFold2). Next, we used the same model, ProstT5, to translate the novel AA sequences back into 3Di sequences that we matched to the starting point (native 3Di or structural template) using percentage pairwise sequence identity (PIDE) applied to 3Di strings (3Di→AA→3Di). If the generated AA sequence adopts the same structure, this will yield high similarity between the source structure (AlphaFold2 prediction of 3Di) and the ProstT5 structure (3Di) prediction for the generated AA sequence. Indeed, this similarity that we dubbed *roundtrip accuracy* correlated well (Spearman R of 0.64) with the structural similarity (lDDT) of 3D predictions of generated and native sequences (Fig. S6A). When giving the model ten attempts to reach a minimal *roundtrip accuracy* (≥70), we observed a minor, yet consistent improvement on all metrics (Table 2).

### Traditional embedding extraction

When merging protein sequences (AA) and structures (3Di) in a single model, we hypothesized such multi-modal pre-training to increase the usefulness of amino acid embeddings as input to subsequent structure-related prediction tasks. Indeed, for secondary structure prediction and the classification of proteins in similar structural classes, ProstT5 embeddings improved over other classification methods with (Fig. 3A, Table S1, Fig. S2A) and without supervision (Table 1). Yet, not all protein prediction tasks benefited directly from coupling 3Di and AA. In fact, tasks related more to function than to structure performed even slightly worse (Fig. 3B-D). For location prediction (Fig. 3C), this might partially be explained by decreased 3D structure prediction precision at the ends of the sequence including the N-terminal signal peptides that are crucial for subcellular localization. For other tasks such as binding or conservation prediction, the drop in prediction performance was less pronounced, i.e., insignificant for conservation, and might be explained by repurposing some of the model’s capacity for structure-specific information. However, we could compensate for this drop by a simple concatenation of the embeddings from the original ProtT5 and the AA-based part of ProstT5 (ProstT5(AA)). In particular, in predicting binding or residues to other ligands (excluding other proteins^17^), the simple concatenation performed best (Fig. S2B: ProtT5 + ProstT5(AA) vs. SOTA in Fig. 3B). Although this increase remained statistically insignificant.

### Limitations

By building a highly non-redundant, and diverse data set consisting only of high-quality 3D structures, we managed to maximize the amount of sequence-structure space covered by a minimum of proteins. While this approach toward reducing redundancy avoided excessive bias towards large families that exists in both the PDB^37^ and in sequence databases^34^, our filter might have introduced another bias. If so, ProstT5 might have amplified bias in its training data like other LLMs^65^. Most important might be bias pertaining to structure predictions and how we filter and represent those (3Di). For instance, filtering the AFDB for predictions with high pLDDT removes most intrinsically disordered proteins^66^ and enriches short^67^, well-structured, preferably helical proteins^68^. On the one hand, this is intended because we want to avoid training on non-structured proteins. On the other hand, the high class imbalance of 3Di-tokens (50% of residues in 10% - 2 of 20 - of the 3Di tokens, Fig. S1), impaired training as some proteins were represented by a single 3Di-token, e.g., most all-helical proteins contained only one 3Di-token (d). We addressed this by removing extreme outliers with more than 95% of the residues being assigned to the same 3Di-token. Nevertheless, class imbalance remained high. Thus, future work might benefit from improved and potentially more balanced 3D→1D mappings.

Another problem might be the circularity and potential information leakage arising from pre-training on 3Di sequences which appear to capture secondary structure well (Fig. 3A -OHE(3Di), Fig. S1) while partly evaluating on secondary structure (Fig. 3A) or hierarchical structure classification (Fig. 2, Table 1). However, this circularity did not impair the practical usefulness of the proposed method as (i) CASP14 performance still improved (Fig. 3A), and (ii) ProstT5 embeddings outperformed one-hot-encoded 3Di (Fig. 3A), as well as, succeeded at detecting CATH homologous superfamilies (Table 1) or SCOPe folds (Fig. 2). The latter required nuanced structural understanding beyond simple secondary structure. As the number of truly novel folds appears to be limited^36,69^, a method contextualizing existing structural motifs well, might be important for many use cases.

### Outlook

LLMs had been phenomenally successful when the release of GPT4 advanced another magnitude. Our proof-of-principle solution rendering protein structures directly amenable to LLMs will benefit from future NLP improvements, in particular, from better sampling of conditional language models^70^. Fine-grained control over this sampling might improve *in silico* creations of predicted multiple sequence alignments (MSAs) when sampling from the ProstT5-improved cycle from single sequence to potentially multiple structure(s) to multiple sequences (AA→3Di**<u>s</u>**→AA**<u>s</u>**). Constant speed-ups of LLMs^60^ make the direct prediction of 3Di from AA an ever more attractive alternative to searching large metagenomic datasets at the sensitivity of structure comparisons, at multiple orders of magnitude of speedup over structure prediction. Along the same lines, predicted 3Di can help speed up recently developed structural phylogenetics based on 3Di^71^. Deriving embeddings from structures will also expand the power of embedding-based alignments^72,73^, and retrieval Transformers^74^. Our proposed integration of 3D information into pLMs constitutes the first step toward building truly multi-model pLMs that capture the multiple facets of protein structure, function and evolution. The first truly bilingual pLM that, in analogy, allows from inputting a text creates an image, from inputting an image provides the subtitles. Expanding ProstT5 into becoming truly polyglot by adding other, potentially more function-centric, conditioning tags such as Gene Ontology terms^75^ might be the next steps toward advancing generic pLMs even more. Recent developments on increasing the context length of LLMs^76^ will enable using full-length UniProt entries, or even all papers that mention a certain protein. This way, LLMs can integrate any knowledge existing for a certain protein today.

## Conclusion

Over the last two years, we have been witnessing how the protein structure revolution ignited by *AlphaFold2* enables groundbreaking scientific discoveries. However, integrating the wealth of information arising from this new, gigantic data resource demands the development of novel tools to optimally leverage the potential. *Foldseek* is a first leap paving the way for new avenues in the era post *AlphaFold2*. ProstT5 exemplifies how language modeling techniques and Transformers can be used for tapping into this information-rich goldmine of 3D structures.

## Availability

Our model is freely available for all at https://huggingface.co/Rostlab/ProstT5 with example scripts deposited at https://github.com/mheinzinger/ProstT5. Additionally, we make PDB files with 3D coordinates available at https://rostlab.org/~deepppi/prosst5_PDBs.tar. 3Di sequences extracted thereof can be queried from https://huggingface.co/datasets/Rostlab/ProstT5Dataset. To make structure search via ProstT5-predicted 3Di sequences available to everyone we also include it into the Foldseek webserver https://search.foldseek.com.

## Methods

### ProstT5 data set

Our translation from 1D amino acid sequences to 1D structure sequences (3Di tokens) began with a recently published^36^ clustered version of the AlphaFold Protein Structure Database (AFDB^33^). This dataset was created by two-step clustering and one step of quality filtering.

(i) MMseqs2^35^ clustered 214 million (M) UniprotKB^34^ protein sequences from AFDB such that no pair had over 50% pairwise sequence identity (PIDE) at 90% sequence overlap. For each of the 52M resulting clusters, the protein with the highest predicted local distance difference test (pLDDT) score^32^ was selected as the representative.
(ii) Foldseek^1^ clustered the 52M representatives further into 18.8M clusters enforcing a pairwise minimal E-value of 10^-2 at 90% sequence overlap for each Foldseek (structural) alignment. From those 18.8M, 2.8M clusters contained two or more members (16M were singletons, i.e., no other protein could be aligned using the procedure above). To avoid bias towards exotic outliers and to increase the training set, we expanded each cluster, maximally, by its 20 most diverse members using HHBlits^77^. This expansion increased from 2.8M clusters to 18.6M proteins leading to a total set size of 34.6M proteins when combined with the singletons.
(iii) Finally, we added three filtering steps: remove (a) low-quality structure predictions (pLDDT<70), (b) short proteins (length<30), and (c) proteins with highly repetitive 3Di-strings (>95% of assigned to single 3Di token). The final training set contained 17M proteins (4.3M singletons with respect to the original 16M). As we are translating into both directions, i.e., from 3Di to amino acids (AA) and vice versa, this corresponded to 34M training samples. From those, we randomly split off 1.2k proteins for validation and 1.2k for final testing while ensuring that clusters were moved to either of the sets such that all members of one cluster always end up in the same split. After keeping only representatives to avoid bias towards clusters, we ended up with 474 proteins for validation and final testing each.

For comparison of the final dataset to PDB^37^ (Fig. S1), we downloaded PDB’s “ss_dis.txt” file (28.06.2023) which simplifies extraction of (un-)resolved residues and their secondary structure elements. Corresponding 3Di sequences were extracted from the PDB version provided by Foldseek. For the analysis, we removed any entry that a) could not be matched between both versions, b) had a length mismatch in the Foldseek 3Di string and the AA sequence length in PDB, c) removed all unresolved residues and d) transformed 8-state secondary structure as defined by DSSP^78^ to 3-states by mapping {G,H,I}→*Helix*, {B,E}→*Strand*, {-,T,S}→*Other*.

### ProstT5 training

#### ProstT5 pre-training

To learn translating between structure (3Di) and sequence (AA), we chose the already pre-trained protein language model (pLM) ProtT5 (marked *ProtT5-XL-U50* in the original publication^6^). We could have started from scratch but wanted to save resources by building on top of existing knowledge. ProtT5 is based on the sequence-to-sequence model T5^38^ trained on reconstructing corrupted tokens from 2.1B metagenomic protein sequences in the BFD (Big Fantastic Database, ^79^) and 40M protein sequences from Uniref50^80^, a version of UniProt^34^ clustered at 50% sequence similarity. We chose this pLM, because the original Transformer consisting of an encoder-decoder architecture that is used by (Prot)T5 lends itself to translation tasks. During training, the encoder learns to parse the source language while the decoder learns to produce meaningful output in the target language conditioned on the encoder output.

#### Learning new 3Di tokens

In a first step, we expanded the existing ProtT5 vocabulary consisting of the 20 standard amino acids and 128 special tokens introduced for span-corruption^38^ by the 20 3Di tokens. To avoid token collision during tokenization of amino acids and 3Di strings (which use identical letters), we cast all 3Di sequences to lower-case before using them as input to the model. Additionally, we added two special tokens (“<FOLD2AA>”, “<AA2FOLD>”) which are prepended to the input to indicate the directionality of the translation. More specifically, *<FOLD2AA>* instructs the model to translate from the input of a 3Di structure sequence into an amino acid sequence, while *<AA2FOLD>* indicates the inverse direction, i.e., instructs the model to generate a 3Di structure sequence from an amino acid input. With this setup, we continued the ProtT5 pre-training on *train17M*, i.e., reconstructing corrupted tokens from non-corrupted context, but now simultaneously using protein sequences (amino acids) and structures (3Di). By training on both modalities simultaneously we tried to avoid catastrophic forgetting which will become important later when translating in both directions. The 3B (3*10^9) free parameters of ProtT5 were fine-tuned with a learning rate of 10^-3 for 10 days on 8 Nvidia A100 each with 80GB vRAM using a batch-size of 16. As pLMs benefit from training on many samples (or manyfold repetitions of the same samples^6^), we increased throughput by limiting sequence lengths to 256 (truncating longer sequences) and using DeepSpeed (stage-2)^81^, gradient accumulation steps (5 steps), mixed half-precision (bf16, ^82^) and PyTorch2.0’s torchInductor compiler^83^. Thereby, the model trained on 102M (1.02*10^8) samples corresponding to about three epochs (1 epoch = presentation of each sample once) over the 34M protein (structure) sequences in *train17M*.

#### Learning bi-directional translation

In a last step, the resulting model, which can now “read” 3Di tokens, was trained to translate between sequences of amino acids and 3Di *structure states*. Both directions were trained simultaneously with the prefixes *<FOLD2AA>* and *<AA2FOLD>* indicating the directionality of the translation. The translation was trained on the same machine (8 A100 á 80GB vRAM) and setup (DeepSpeed stage-2, gradient accumulation, bf16, torchInductor) as before. However, we changed the learning rate to 10^-5 for the initial 100K (10^5) steps (6 days) on sequences with ≤256 residues (again truncating longer sequences), and increased to 512 for another 600K (6*10^5) steps (20 days). While increasing sequence length, we had to lower batch-size (from 16 to 6) which we compensated for by increasing the number of gradient accumulation steps (from 1 to 4). In total, we trained for around 700K (7*10^5) steps (about 4 epochs) on set *train17M*. We dubbed the final model *<u>ProstT5</u>*for *<u>Protein structure sequence T5</u>*.

### Evaluation benchmarks

#### Transfer learning

One way to establish the value of pLMs is to use the vector representations they learned, referred to as the embeddings, as input to subsequent supervised prediction tasks^5^. Ultimately, this is the concept of transfer learning which, in the first step, requires substantial computing resources to create general purpose pLMs. In the second step, the embeddings from these pLMs, i.e. the essence of what they learnt, are input to any supervised prediction task of interest. In this logic, the performance of some standardized, non-optimized set of 2^nd^ step supervised prediction tasks becomes the best way to evaluate the validity of the pLM (here ProstT5) as a general-purpose model. As redundancy between training and testing is crucial even in relative comparisons^84^, this aspect makes it so difficult to adequately evaluate protein prediction on known data^43,85^, which is why we focused on a limited number of standard benchmarks that we tried to reproduce as closely as possible to existing work using biotrainer^41^ and FLIP^42^.

#### Supervised learning: per-residue: secondary structure

Given its proven track record to benchmark pLMs^5–8,12^ and to ease comparison to other methods, we replicated previous benchmarks^6^. To predict properties of single tokens (here: single amino acids, dubbed residues when joined in proteins), we used the training set published with NetSurfP-2.0^46^ (3-state secondary structure using DSSP^78^: helix, strand, and other). We benchmarked using three public test data sets, namely CASP12^43^, CASP14^44^ and NEW364^6^. We report performance on CASP12 and NEW364 for comparability to existing methods but those sets allow for indirect information leakage as they overlap with AlphaFold2 training data (and thus with our set *train17M*). We used the same convolutional neural network and hyperparameters as described in detail elsewhere^6^.

#### Supervised learning: per-residue: binding

For predicting whether a ligand (small molecule, metal ion, or DNA/RNA; essentially only excluding protein-protein interactions) is binding to a specific residue in a protein, we replicated training (*DevSet1014*) and testing (*TestSet300*) of a recent method^17^ (also using a two-layer CNN; with the same training parameters). For simplicity, we skipped the more fine-grained evaluation of different binding types focusing on the binary binding/not.

#### Supervised learning: per-residue: conservation

One surprising recent result established that pLMs can reliably predict the conservation of a residue in a protein family without using any multiple sequence alignment defining a family as input^45^. Here, we replicated the training and evaluation used before^45^. In brief, we used *ConSurf10k*^45,86^, a 25% non-redundant dataset derived from high-quality PDB^37^ structures, to train a two-layer CNN to classify each residue into one of nine conservation classes (1=highly variable, 9=highly conserved) defined by ConSurf-DB^86^.

#### Supervised learning: per-residue: 3Di classification

To speed-up the search for related proteins based on ProstT5-generated 3Di sequences, we dropped ProstT5’s decoder and instead trained a two-layer CNN on top of embeddings extracted from the ProstT5 encoder to classify each residue in a protein into one of the 20 3Di states. For training, we used the same network architecture, hyperparameters, sets and splits as for secondary structure prediction. Instead of deriving 3Di-states directly from experimental data, we trained on 3Di-states extracted from AlphaFold2 predictions to avoid inconsistencies such as missing or mutated residues part of experimental PDB files.

#### Supervised learning: per-protein: subcellular location

For predicting features of entire proteins, we classified each protein into one of ten subcellular locations^18^. More specifically, we used the *DeepLoc* training data^87^ to train a light-attention network^18^ which we evaluated using a 20% non-redundant test set (*setHARD*). We copied setup and hyperparameters from the literature^18^.

#### Supervised learning: per-protein: superfamily classification

We used CATH^50^ to classify proteins into superfamilies replicating previous work ^21^. In brief, we used the CATH hierarchy (v4.3) which classifies three-dimensional (3D) protein structures at the four levels Class (most distantly related pairs), Architecture, Topology and Homologous superfamily (most similar pairs). As described in ^21^, we used contrastive learning to train a two-layer feed-forward-neural network to learn a new embedding space in which proteins with increasing overlap in the hierarchy are pulled closer together while others get pushed further away. A hit at lower CATH-levels could be correct if all previous levels were correctly predicted. Due to the varying number of samples at different CATH-levels, performance measures not normalized by background numbers could be higher for lower levels.

#### Unsupervised classification: per-protein: superfamily prediction

Other than serving as input to 2^nd^-step supervised training, embeddings can also be used without further modifications for classifications directly. One solution is the so-called embedding-based annotation transfer (<u>EAT</u> - ^14,21^) that proceeds as follows. Given a protein K of experimentally known annotation and another protein Q with missing annotation: if the Euclidean distance between the two is below some empirical threshold T (if D(embedding(Q),embedding(K))<T): transfer annotation of K to Q. Arguably, EAT generalizes what is used for most database annotations and is often referred to as homology-based inference (HBI) that copies annotations when Q is sufficiently sequence similar to K. In order to classify/predict, the annotation of the most similar protein in the lookup database is transferred to the query protein.

We used a previously published dataset^21^ to probe how well ProstT5 per-protein embeddings (average-pooling over the residue-embeddings derived fixed-length vector for each protein irrespective of its length) alone distinguished between different levels of structural similarity. Instead of applying supervision and contrastive learning, we used EAT to transfer annotations from a lookup set to a 20% non-redundant test set. For protein sequences, this task corresponded to something as daring as using 20d-vectors with the amino acid composition of two proteins to establish whether or not those have similar structure. Again, we computed the accuracy as the fraction of correct hits for each CATH-level.

ProstT5’s translation capabilities also open other new possibilities for unsupervised benchmarking of the information stored in the model. For example, one can use ProstT5 to translate from sequence to structure and use predicted structure (3Di) for remote homology detection.

#### *Folding*: from sequence to structure

An alternative way to extract information from ProstT5 is to predict 3Di sequences from amino acid sequences (Fig. 1) and use the predicted 3Di sequences as input to *Foldseek* to search for (structurally) related proteins. Towards this end, we reproduced the *Foldseek* benchmark replacing 3Di strings derived from experimental data by ProstT5 predictions. In brief, Foldseek performs an all-against-all search of SCOPe40 (SCOPe 2.01 clustered at 40% pairwise sequence identity^39^) to measure the fraction of finding members of the same SCOPe family, superfamily and fold (true-positive (TP) matches) for each query out of all possible correct matches until the first false positive (FP: match to different fold). *Foldseek*, when used on PDB structures, uses C-alpha backbone information to rerank hits, which slightly improves performance in the SCOPe40 benchmark. Since no C-alpha information is available when using ProstT5 to generate 3Di strings, we disabled this feature to evaluate ProstT5 but activated it when running Foldseek for fair comparison.

#### *Inverse folding*: from structure to sequence

The term *inverse folding*^88,89^ has been applied to the challenge of finding all the protein sequences that adopt a particular 3D structure (in the past loosely referred to as “the fold”). By design ProstT5 appears ideally suited to address this challenge by simply inverting the direction of the translation, i.e., by reconstructing amino acid sequences from 3Di-sequences. Toward this end, we considered sequence similarity to be a weak measure for success as there are many, potentially very dissimilar, sequences that still adopt the same structure. Instead, we used structural similarity, i.e., Local Distance Difference Test (lDDT^57^), template-modeling score (TM-score^58^) and root-mean-square-deviation (RMSD). To obtain 3D coordinates, we predicted structures using ESMFold^8^ for all protein sequences created by ProstT5 and compared these predictions to the AFDB *groundtruth* for the native sequence.

#### Sampling from translations

In contrast to traditional classification, so-called conditional language models assign probabilities to a sequence of words given some conditioning context. Here, we either generated amino acid sequences conditioned upon structure (3Di sequences) or vice versa. As there are multiple techniques to sample from this process, each with individual hyperparameters, we compared different sampling strategies^53–55^. All comparisons and resulting decisions were based on the validation set while the final testing set was only used to report final performance of the hyperparameter combination that worked best on the validation set.

*AA→3Di (folding):* When translating from amino acid (AA) sequences to 3Di-sequences, we used the sequence similarity (below) between the groundtruth 3Di sequences from the AFDB and the ESMFold predictions from the generated 3Di sequences to compare different sampling strategies. We used global Needleman-Wunsch alignment^90^ as implemented in biotite^91^ together with the 3Di substitution matrix from Foldseek^1^ to compute sequence similarities. We compared all combinations of the following parameters (SOM: Table S2): a) number of beams ∈ [0,3] ^53^, b) temperature ∈ [0.8, 1.0, 1.2], c) top-p ∈ [0.85, 0.9, 0.95] ^54^, d) top-k ∈ [3,6] ^55^, and, e) repetition penalty ∈ [1.0, 1.2, 1.4]. For all analysis presented here, we used the following huggingface generation configuration because it achieved the highest sequence similarity: "do_sample": True, "num_beams": 3, "top_p" : 0.95, "temperature" : 1.2, "top_k" : 6, "repetition_penalty" : 1.2. *3Di→AA (inverse folding):* To reconstruct amino acid sequences (AA) from 3Di-sequences, we again compared all combinations of the following sampling parameters (SOM: Table S3): a) number of beams ∈ [0,3], b) temperature ∈ [0.8, 0.9, 1.0, 1.1, 1.2], c) top-p ∈ [0.85, 0.9, 0.95], d) top-k ∈ [3,6], and, e) repetition penalty ∈ [1.0, 1.2, 1.4]. However, this time, we defined success in terms of a combination of the lDDT (comparing ESMFold predictions of our generated sequences against AFDB) and naturalness as proxied by relative entropy (or Kullback-Leibler divergence) between the amino acid distribution in UniProt and the generated sequences^56^. This resulted in the following configuration: "do_sample": True, "top_p" : 0.85, "temperature" : 1.0, "top_k" : 3, "repetition_penalty" : 1.2.

#### Proteome runtime benchmark

We compare the 3D structure prediction runtime of the 48h shown in the ColabFold manuscript using an optimized prediction workflow to 3Di string prediction using the ProstT5 encoder-CNN mode. Here, we take the same *M. jannaschii* proteome (UniProt proteome accession UP000000805) subset of all sequences shorter than 1000 AA and predict the runtime on a Macbook Pro 13” M1 2020 with 16GB RAM and report the average run-time and standard deviation of five repeated runs. The commandcall executed was hyperfine^92^ --runs 5 ’python3 predict_3Di_encoderOnly.py -i UP000000805_243232_lt1000.fasta -o UP000000805_243232_lt1000_3di --model model --half 0 --output_probs 0’. We repeated the same procedure on a server with 1x AMD EPYC 7402P, 512GB RAM and 8x Nvidia RTX A5000, of which one was utilized. Additionally we enabled the half precision mode for further speedup (--half 1), which was disabled for CPU as it is currently unsupported and results in crashes.

## Supporting information

Supplementary Online Material (SOM)

## Acknowledgements

Thanks primarily to the team at the Leibniz Supercomputing Center (LRZ, Munich), especially to Juan Durillo Barrionuevo, for providing access and guidance which enabled large-scale GPU training. Thanks to Chris Dallago (Nvidia), Adam Grzywaczewski (Nvidia) and Noelia Ferruz (IBMB, Spain) for helpful discussions. Thanks to Tim Karl (TUM) for invaluable help with hardware and software and to Nicola Bordin (UCL) for providing access to the non-redundant PDB structures of CATH. MH and BR were supported by the Bavarian Ministry of Education through funding to the TUM, by a grant from the Alexander von Humboldt foundation through the German Ministry for Research and Education (BMBF: Bundesministerium für Bildung und Forschung), and by a grant from Deutsche Forschungsgemeinschaft (DFG-GZ: RO1320/4-1). MS acknowledges the support by the National Research Foundation of Korea, grants [2020M3-A9G7-103933, 2021-R1C1-C102065, 2021-M3A9-I4021220], Samsung DS research fund and the Creative-Pioneering Researchers Program through Seoul National University. MM acknowledges support by the National Research Foundation of Korea (grant RS-2023-00250470). Last, but not least, thanks to all those who maintain public databases, in particular, to the team at EMBL-EBI who teamed-up with Deepmind to make AlphaFold2 3D structure predictions for hundreds of millions of proteins in UniProt publicly available to everyone.

## Abbreviations & Glossary

1D: one-dimensional (string such as secondary structure)
3D: three-dimensional (coordinates)
3Di: 1D-strings representing protein 3D structure (taken from Foldseek^1^)
AA: amino acid
AFDB: AlphaFold Protein Structure Database
CATH: hierarchical classification of protein 3D structures in Class, Architecture, Topology and Homologous superfamily
CNN: convolutional neural network
EAT: Embedding-based Annotation Transfer; embeddings fixed-size vectors derived from pre-trained pLMs
LLM: large language model
pLM: protein Language Model
SOTA: state-of-the-art.

