## Supplementary Online Material (SOM) for "Bilingual Language Model for Protein Sequence and Structure"

In this Supporting Online Material (SOM), we provide more information on the performance of refining ProstT5 embeddings for CATH[1] fold classification using contrastive learning (Table S1). In Fig. S1 we provide more details on dataset statistics, most importantly on how certain 3Di tokens are biased towards specific secondary structure states (Fig. S1B). Additionally, we give more details on the effect of concatenating various embedding types before using them for supervised training of various downstream tasks (Fig. S2). In Fig. S3 we compare different ways to predict 3Di from amino acid sequence only. The prediction mistakes of the best performing model from this evaluation are then analysed in more detail via a confusion matrix (Fig. S4) and the effect of varying output probability thresholds on precision and recall are measured (Fig. S5). In Fig. S6 we further highlight how the self-consistency, i.e., translating from 3Di→AA→3Di and comparing similarity of start/end sequences, correlates with the quality of inverse folding quality.
Finally, we provide details on the generation configurations that we compared for translating from amino acid sequences to 3Di-sequences (Table S2) and vice versa (Table S3).

**Table S1: ProstT5 embeddings for fold classification (contrastive learning).** **2**
**Figure S1: Dataset analysis.** **3**
**Figure S2: Protein prediction tasks using concatenations of pLM embeddings.** **4**
**Figure S3: Comparison of different approaches for predicting 3Di.** **5**
**Figure S4: Confusion matrices for 3Di prediction from ProstT5(AA)+CNN.** **6**
**Figure S5: Precision and coverage tradeoff for 3Di predictions.** **7**
**Figure S6: ProstT5 back-translation correlates with inverse folding quality.** **8**
**Table S2: Generation parameters for translating sequence (AA) to structure (3Di).*** **9**
**Table S3: Generation parameters for translating structure (3Di) to sequence (AA).*** **11**

**Table S1: ProstT5 embeddings for fold classification (contrastive learning).**

| | Method/Input | C | A | T | H | Mean |
|---|---|---|---|---|---|---|
| EAT - supervised | ESM-1b‡ | 87±4 | 68±6 | 59±7 | 70±7 | 71±6 |
| | Ankh | 87±4 | 76±6 | 64±6 | 75±7 | 75±6 |
| | ProtT5‡ | 89±4 | 75±6 | 64±6 | 76±6 | 76±6 |
| | ProstT5(AA) | 89±4 | **81±5** | 70±6 | 83±6 | **81±5** |
| | ProstT5(p3Di) | 88±4 | 77±5 | 69±6 | 80±6 | 79±5 |
| | ProstT5(3Di) | 90±4 | 78±5 | 69±6 | 74±7 | 78±6 |
| | ProstT5(cat) | **91±4** | 79±5 | **72±6** | **84±6** | **81±5** |

*Accuracy for predicting CATH [1] levels (from coarse- to fine-grained: C, A, T, H) by transferring annotations from a lookup set to a strictly non-redundant set of queries. The column *Mean* marked the simple arithmetic average over the four performance values. Values of methods marked with ‡ are taken from [2]. Methods: EAT: supervised: contrastive learning optimised on CATH for input embeddings from ESM-1b[3] , Ankh [4], ProtT5 [5] and ProstT5 (same inputs as for unsupervised). ProstT5(cat) marks the concatenation of ProstT5(AA) and ProstT5(p3Di).

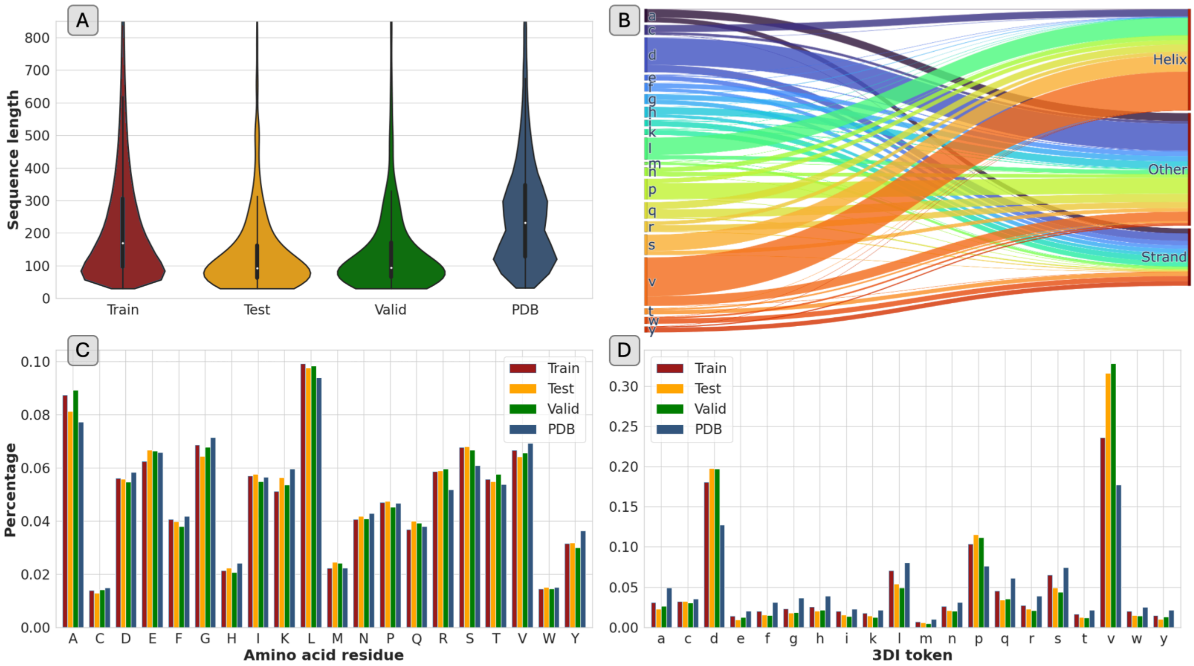


**Figure S1: Dataset analysis.**

We compared basic properties (A: sequence length, C: 3Di-distribution, and, D: amino acid composition) of the proteins in our data sets (blue: training, green: validation, orange: test set) with experimentally resolved proteins (dark blue: PDB[6]). The PDB data also revealed the relation between 3Di-tokens (panel B: left: 3Di states in lower-case for distinction to amino acids; right: secondary structure states, i.e., helix, strand and other). While the amino acid distribution was similar between predicted (AFDB[7]) and experimental (PDB) structures (D), some 3Di-tokens (d, v, p) were clearly more over-represented in AlphaFold2 predictions than in the PDB. The Sankey diagram (B), revealed those tokens mostly related to helix (3Di: v) or other (loop; 3Di: d and p). In contrast, 3Di tokens accounting mostly for strands (a, w, y) appeared more frequently in the PDB than in our data. Proteins in our data also tended to be slightly shorter than proteins in PDB, with average lengths of 206-238 (test-train) and 255, respectively.

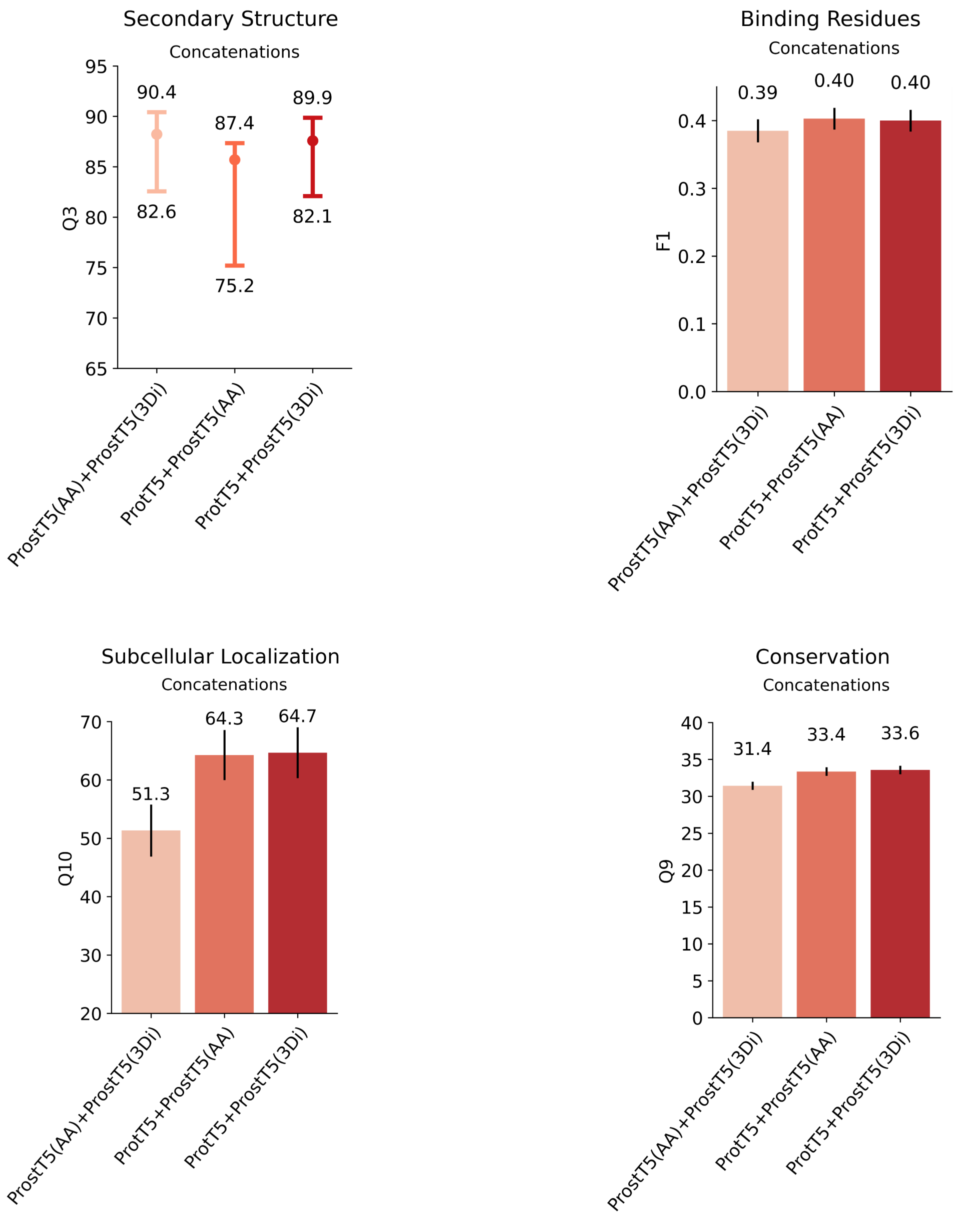


**Figure S2: Protein prediction tasks using concatenations of pLM embeddings.**

We probed whether different embedding sources provide orthogonal information that might improve over each individually by concatenating embeddings from sequences (ProstT5(AA) and ProtT5) and structures (ProstT5(3Di)) and using them as input to subsequent supervised prediction methods. We again compared four different prediction tasks, namely the per-residue prediction of secondary structure (A; performance: Q3: three-state per-residue accuracy; data sets: middle: CASP12[8], lower bar: CASP14[9], upper bar: NEW364[5]; note: since each of those sets is supposed to measure performance, the difference between them proxied the error in the estimate), binding residues (B; performance: F1; data: testSet300[10]), conservation (D; performance: Q9: nine-state per-residue accuracy; data: [11]), and the per-protein prediction of subcellular location (C; performance: ten-state per-protein accuracy, Q10; data: setHARD[12]). For panels B-D, the bars mark the 95% confidence interval, i.e., ±1.96 * standard errors, estimated via bootstrapping.

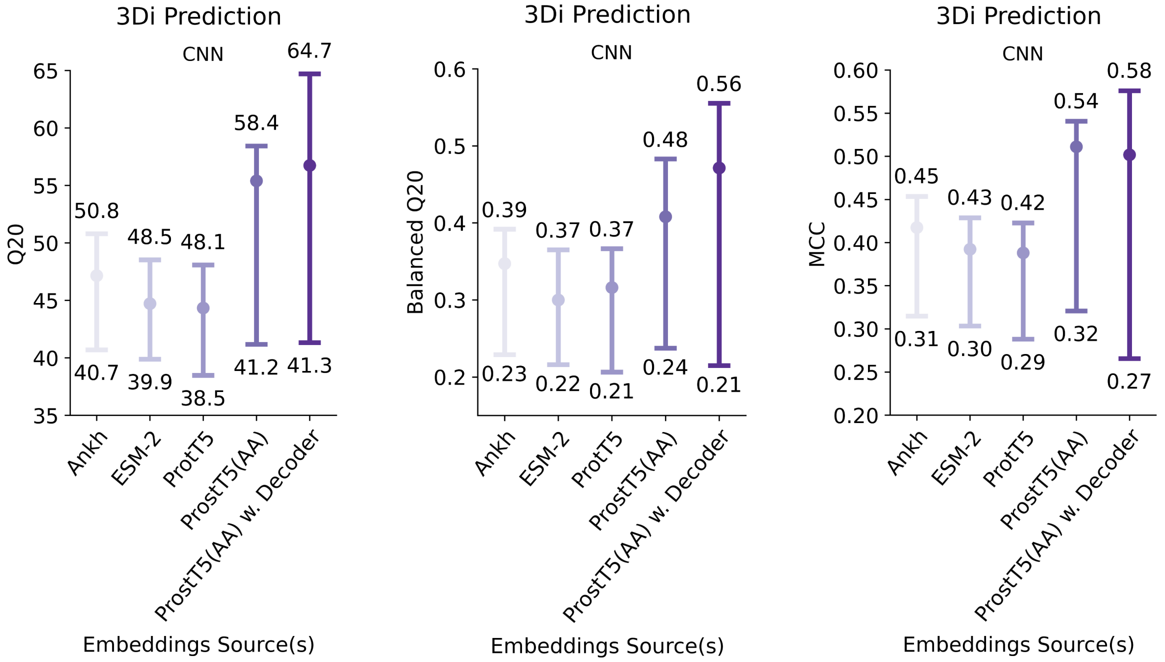


**Figure S3: Comparison of different approaches for predicting 3Di.**

We compared different approaches and embedding sources for predicting 3Di directly from amino acid sequences. Ground truth 3Di states were derived from AlphaFold2 (AF2) predictions for the secondary structure sets used throughout this work (middle: CASP12[8], lower bar: CASP14[9], upper bar: NEW364[5]). Using AF2 predictions instead of experimental structures bypassed mapping issues originating from missing or unresolved atoms in experimental structures. We compared training a 2-layer CNN on top of embeddings derived from either Ankh[4], ESM-2 (3B)[13], ProtT5[5] or ProstT5's encoder (ProstT5(AA)) against using ProstT5's encoder-decoder (ProstT5(AA) w. Decoder) to predict 3Di sequences. In all cases, we only used amino acid sequences as input. To account for the class imbalance of 3Di-tokens (Fig. S1D), we compared 20-state classification accuracy (Q20) against balanced accuracy (balanced Q20) and Matthews-correlation-coefficient (MCC[14]) which were tailored for handling class imbalance. The latter comparison revealed an ambiguous trend: while the ProstT5 decoder was better for NEW364, the CNN trained on top of ProstT5's encoder was better on the arguably hardest set, CASP14 (lowest bar), and was on par for CASP12.

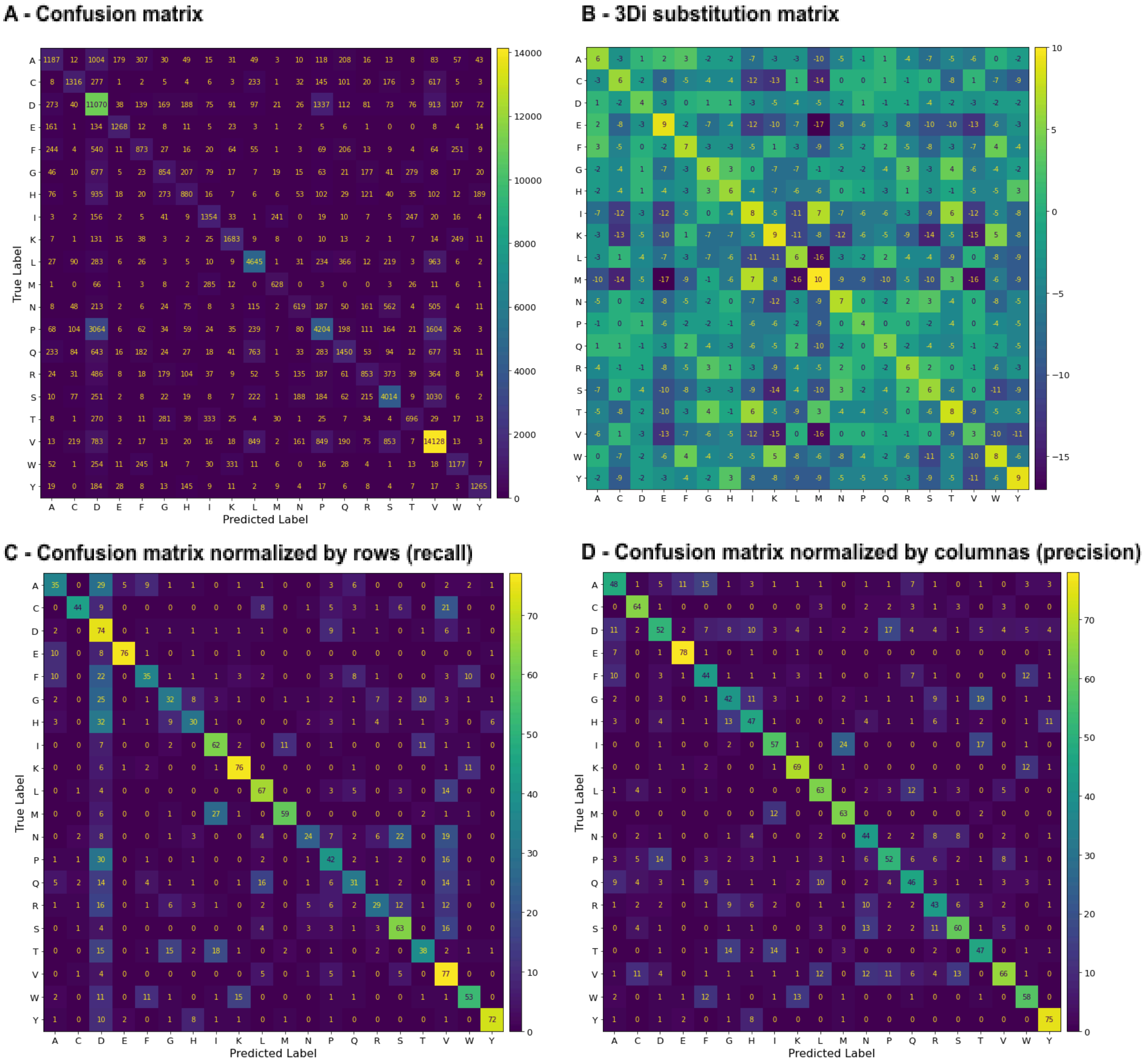


**Figure S4: Confusion matrices for 3Di prediction from ProstT5(AA)+CNN.**

To better understand prediction mistakes made by the CNN trained on top of embeddings derived from inputting amino acids to the encoder of ProstT5 (ProstT5(AA)), we visualised the raw confusion matrix (Panel A) comprising all three test sets (CASP12[8], CASP14[9], and NEW364[5]) together with its normalised counterparts (Panel C showing normalisation over true labels or rows, i.e. recall, and Panel D showing normalisation over predictions or columns, i.e.,precision). For comparison, we also add Foldseek's 3Di substitution matrix (Panel B). Many prediction mistakes, e.g., "P" being confused for a "D", happen amongst 3Di states likely to substitute each other (positive substitution score). Recall for over-represented classes, e.g., "D" and "V", is comparatively high which does not get mirrored in the corresponding precision entries indicating some level of overprediction for the most abundant classes.

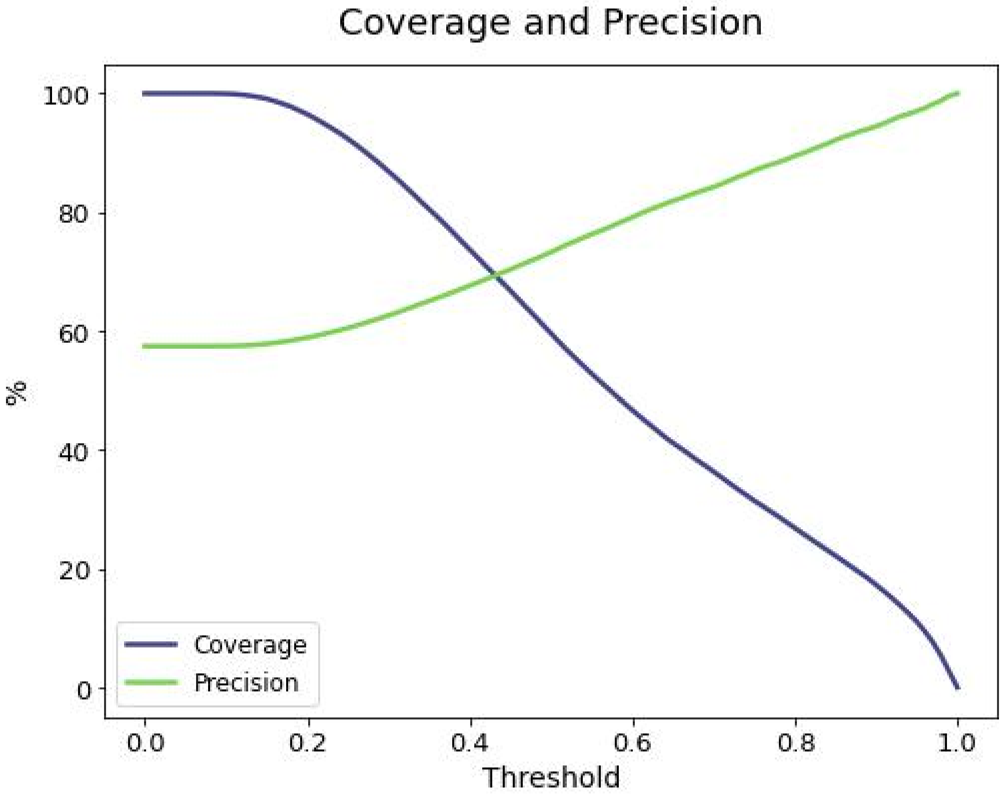


**Figure S5: Precision and coverage tradeoff for 3Di predictions.**

We further investigated the tradeoff between coverage (blue) and precision (green) when applying different thresholds to the output probabilities of the CNN trained on top of ProstT5's encoder to predict 3Di states from amino acid embeddings. For example, when only considering predictions with an output probability of 0.5 or better, the proposed method can predict 70% of the residues at 70% precision. With a more conservative threshold of 0.8, the method predicts only for about 25% of the residues, albeit at 90% precision.

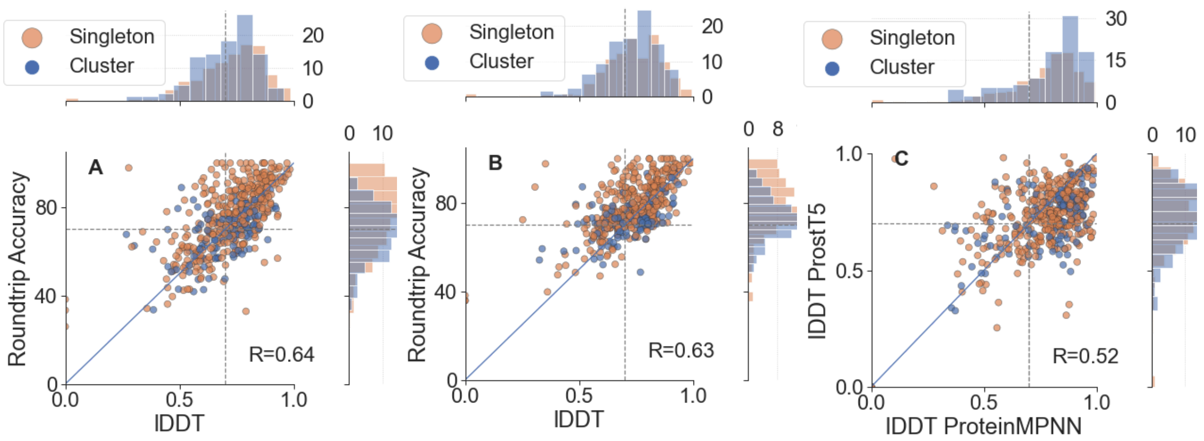


**Figure S6: ProstT5 back-translation correlates with inverse folding quality.**

For each protein in our test set, we used ProstT5 to generate new AA sequences from its native counterpart described by its 3Di sequence (3Di→AA) with the constraint that newly generated sequences share similar structures (measured by lDDT [15]). For the native structures, we took AlphaFold2[16] predictions. For the newly generated sequences, we used single-sequence-based ESMFold[13] to avoid bias towards multiple sequence alignments (MSAs). We dubbed the similarity between 3Di sequences derived from native structures and the 3Di sequences predicted by ProstT5 from the newly generated proteins “Roundtrip accuracy” ((3Di→AA→3Di, y-axis in A and B). A perfect roundtrip recovers the difference between ESMFold and AlphaFold2, i.e, is below 100% PIDE (in terms of 3Di). Panel A plots the Roundtrip accuracy against the structural similarity (lDDT) between ESMFold predictions for mutant sequences and AlphaFold2 for the native. Panel B leveraged this correlation (R=0.64) to let ProstT5 control its mutations by generating AA sequences until either a Roundtrip accuracy≥70 was reached or for maximally ten attempts. This constraint slightly decreased the Pearson R (0.64 vs. 0.63) but improved the structural similarity between the newly generated sequences (Table 2). Panel C compared the same ProstT5 generations (constrained by roundtrip accuracy, panel B) with ProteinMPNN[17] in terms of lDDT. There seems to be a common trend towards easy inverse folding targets as both methods agreed well in terms of lDDT (Spearman R of 0.52). In all cases we differentiated between test proteins in clusters, (blue, clusters being proteins that shared sequence or structure similarity according to data preparation described in Methods) and singletons (orange). Margins: distribution of both classes as percentage values. Dashed lines mark a lDDT or roundtrip accuracy of 70.

Table S2: Generation parameters for translating sequence (AA) to structure (3Di).*

| beam-search | p-value | temperature | k-value | repetition-penalty | median | mean |
|---|---|---|---|---|---|---|
| **b3** | **p0.95** | **t1.20** | **k6.0** | **r1.2** | **75** | **71.746624** |
| b3 | p0.90 | t1.00 | k6.0 | r1.2 | 74.8 | 71.540084 |
| b3 | p0.95 | t1.00 | k3.0 | r1.4 | 74.7 | 71.364346 |
| b3 | p0.85 | t1.00 | k6.0 | r1.2 | 74.7 | 71.454008 |
| b3 | p0.95 | t1.00 | k3.0 | r1.0 | 74.7 | 71.428903 |
| b3 | p0.85 | t1.00 | k3.0 | r1.2 | 74.65 | 71.367932 |
| b3 | p0.85 | t1.20 | k3.0 | r1.2 | 74.65 | 71.466456 |
| b3 | p0.95 | t1.00 | k6.0 | r1.0 | 74.65 | 71.383544 |
| b3 | p0.90 | t1.20 | k3.0 | r1.2 | 74.6 | 71.374262 |
| b3 | p0.85 | t1.00 | k6.0 | r1.0 | 74.55 | 71.344937 |
| b3 | p0.95 | t1.20 | k3.0 | r1.4 | 74.55 | 71.516034 |
| b3 | p0.90 | t1.20 | k3.0 | r1.4 | 74.55 | 71.68903 |
| b3 | p0.90 | t1.00 | k6.0 | r1.4 | 74.55 | 71.594093 |
| b3 | p0.90 | t1.20 | k6.0 | r1.4 | 74.55 | 71.602743 |
| b3 | p0.95 | t1.20 | k3.0 | r1.2 | 74.55 | 71.531646 |
| b3 | p0.90 | t1.00 | k6.0 | r1.0 | 74.5 | 71.391772 |
| b3 | p0.95 | t1.00 | k6.0 | r1.2 | 74.5 | 71.630169 |
| b3 | p0.85 | t1.20 | k6.0 | r1.2 | 74.45 | 71.56962 |
| b3 | p0.90 | t1.00 | k3.0 | r1.0 | 74.45 | 71.205907 |
| b3 | p0.95 | t1.20 | k6.0 | r1.4 | 74.45 | 71.567511 |
| b3 | p0.90 | t1.00 | k3.0 | r1.4 | 74.45 | 71.473418 |
| b3 | p0.95 | t1.00 | k3.0 | r1.2 | 74.4 | 71.466667 |
| b3 | p0.95 | t1.00 | k6.0 | r1.4 | 74.4 | 71.650422 |
| b3 | p0.90 | t1.20 | k6.0 | r1.2 | 74.4 | 71.74557 |
| b3 | p0.85 | t1.00 | k3.0 | r1.4 | 74.35 | 71.449789 |
| b3 | p0.95 | t1.20 | k3.0 | r1.0 | 74.35 | 71.07616 |
| b3 | p0.85 | t1.00 | k3.0 | r1.0 | 74.35 | 71.065823 |
| b3 | p0.85 | t1.00 | k6.0 | r1.4 | 74.35 | 71.247468 |
| b3 | p0.95 | t1.20 | k6.0 | r1.0 | 74.35 | 71.533755 |
| b3 | p0.90 | t1.20 | k3.0 | r1.0 | 74.3 | 71.416667 |
| b3 | p0.85 | t1.20 | k3.0 | r1.4 | 74.25 | 71.596835 |
| b3 | p0.85 | t1.20 | k6.0 | r1.4 | 74.2 | 71.651477 |
| b3 | p0.90 | t1.00 | k3.0 | r1.2 | 74.15 | 71.42173 |
| b3 | p0.85 | t1.20 | k6.0 | r1.0 | 74.05 | 71.08692 |
| b3 | p0.90 | t1.20 | k6.0 | r1.0 | 73.95 | 71.170675 |
| greedy | | | | | 73.95 | 71.117722 |
| b3 | p0.85 | t1.20 | k3.0 | r1.0 | 73.85 | 71.516878 |
| b0 | p0.90 | t0.80 | k3.0 | r1.0 | 72.6 | 70.139451 |
| b0 | p0.85 | t0.80 | k3.0 | r1.0 | 72.5 | 70.272996 |
| b0 | p0.95 | t0.80 | k6.0 | r1.0 | 72.05 | 69.788608 |
| b0 | p0.90 | t0.80 | k6.0 | r1.0 | 72.05 | 70.109705 |
| b0 | p0.85 | t0.80 | k6.0 | r1.0 | 71.75 | 69.927215 |
| b0 | p0.85 | t1.00 | k3.0 | r1.0 | 71 | 69.790084 |
| b0 | p0.85 | t1.20 | k3.0 | r1.0 | 70.8 | 68.595992 |
| b0 | p0.90 | t1.00 | k3.0 | r1.0 | 70.8 | 69.279114 |
| b0 | p0.95 | t1.00 | k3.0 | r1.0 | 70.75 | 68.498734 |
| b0 | p0.95 | t0.80 | k3.0 | r1.0 | 70.7 | 69.598734 |
| b0 | p0.85 | t1.00 | k6.0 | r1.0 | 70.65 | 69.321941 |
| b0 | p0.85 | t1.20 | k6.0 | r1.0 | 70.6 | 68.244515 |
| b0 | p0.90 | t1.00 | k6.0 | r1.0 | 70.2 | 68.624262 |
| b0 | p0.95 | t1.00 | k6.0 | r1.0 | 69.6 | 67.708017 |

| | | | | | | |
|---|---|---|---|---|---|---|
| b0 | p0.90 | t1.20 | k3.0 | r1.0 | 69.1 | 67.995992 |
| b0 | p0.85 | t0.80 | k3.0 | r1.2 | 68.45 | 67.147257 |
| b0 | p0.95 | t1.20 | k6.0 | r1.0 | 68.4 | 65.942616 |
| b0 | p0.90 | t1.20 | k6.0 | r1.0 | 68.35 | 66.794726 |
| b0 | p0.95 | t1.20 | k3.0 | r1.0 | 68.25 | 67.250211 |
| b0 | p0.85 | t0.80 | k6.0 | r1.2 | 67.55 | 65.993038 |
| b0 | p0.90 | t0.80 | k6.0 | r1.2 | 67.4 | 65.937553 |
| b0 | p0.90 | t0.80 | k3.0 | r1.2 | 67.3 | 66.300211 |
| b0 | p0.85 | t1.00 | k3.0 | r1.2 | 66.95 | 65.768776 |
| b0 | p0.95 | t0.80 | k3.0 | r1.2 | 66.7 | 66.187342 |
| b0 | p0.95 | t0.80 | k6.0 | r1.2 | 66.35 | 64.967932 |
| b0 | p0.85 | t1.00 | k6.0 | r1.2 | 65.8 | 64.493671 |
| b0 | p0.90 | t1.00 | k3.0 | r1.2 | 65.65 | 64.663291 |
| b0 | p0.90 | t1.00 | k6.0 | r1.2 | 65.3 | 63.52173 |
| b0 | p0.85 | t1.20 | k3.0 | r1.2 | 65.3 | 64.411392 |
| b0 | p0.95 | t1.00 | k3.0 | r1.2 | 64.75 | 63.842827 |
| b0 | p0.90 | t1.20 | k3.0 | r1.2 | 64.7 | 63.496624 |
| b0 | p0.85 | t1.20 | k6.0 | r1.2 | 63.9 | 63.040295 |
| b0 | p0.95 | t1.00 | k6.0 | r1.2 | 63.9 | 61.847046 |
| b0 | p0.90 | t0.80 | k3.0 | r1.4 | 63.75 | 61.98692 |
| b0 | p0.85 | t0.80 | k3.0 | r1.4 | 63.45 | 62.322996 |
| b0 | p0.85 | t0.80 | k6.0 | r1.4 | 62.8 | 61.707806 |
| b0 | p0.90 | t1.20 | k6.0 | r1.2 | 62.6 | 61.227426 |
| b0 | p0.95 | t1.20 | k3.0 | r1.2 | 62.1 | 61.775738 |
| b0 | p0.85 | t1.00 | k3.0 | r1.4 | 62 | 60.92827 |
| b0 | p0.95 | t0.80 | k6.0 | r1.4 | 62 | 60.071097 |
| b0 | p0.90 | t0.80 | k6.0 | r1.4 | 61.9 | 60.718565 |
| b0 | p0.95 | t0.80 | k3.0 | r1.4 | 61.05 | 60.555485 |
| b0 | p0.90 | t1.00 | k3.0 | r1.4 | 60.4 | 59.945992 |
| b0 | p0.95 | t1.20 | k6.0 | r1.2 | 60.4 | 59.433755 |
| b0 | p0.95 | t1.00 | k3.0 | r1.4 | 60 | 58.942194 |
| b0 | p0.85 | t1.00 | k6.0 | r1.4 | 59.9 | 58.976793 |
| b0 | p0.85 | t1.20 | k3.0 | r1.4 | 59.8 | 58.876793 |
| b0 | p0.90 | t1.00 | k6.0 | r1.4 | 58.9 | 57.947468 |
| b0 | p0.90 | t1.20 | k3.0 | r1.4 | 58.4 | 57.646624 |
| b0 | p0.85 | t1.20 | k6.0 | r1.4 | 57.25 | 56.698734 |
| b0 | p0.95 | t1.20 | k3.0 | r1.4 | 56.55 | 55.70654 |
| b0 | p0.95 | t1.00 | k6.0 | r1.4 | 55.95 | 56.100844 |
| b0 | p0.90 | t1.20 | k6.0 | r1.4 | 55.65 | 54.631646 |
| b0 | p0.95 | t1.20 | k6.0 | r1.4 | 52.9 | 52.349367 |

* Sequence similarity for different generation configurations tested for translating from amino acid sequences to 3Di sequences for proteins in the validation set. Sequence similarity was computed between *groundtruth* (3Di derived from AlphaFold2 predictions[16]) and *predictions* (ProstT5: AA→3Di) using Foldseek's[18] 3Di-specific substitution matrix and global Needleman-Wunsch alignment[19]. The columns show a) the number of beams (b0=no beam, b3=3 beams), b) the p-value (p0.9 means p=0.9), c) the temperature (t0.80 refers to temperature=0.8), d) the k-value (k3.0 means k-value=3), e) the repetition penalty (r1.2 means repetition-penalty=1.2), f) and g) the median and mean sequence similarity reached for a given generation configuration, respectively. For comparison, we also added the performance of simple greedy sampling (always picking the next token based on highest probability). For the ProstT5 paper we chose the configuration highlighted in bold.

Table S3: Generation parameters for translating structure (3Di) to sequence (AA).*

| beam | p-value | temperature | k-value | repetition-p | lDDT | Entropy | TM-score | PIDENT |
|---|---|---|---|---|---|---|---|---|
| b3 | p0.85 | t1.10 | k3.0 | r1.2 | 0.77375 | 0.089946 | 0.59 | 22.2 |
| b3 | p0.85 | t1.30 | k3.0 | r1.0 | 0.77145 | 0.101324 | 0.59 | 21.9 |
| b3 | p0.95 | t1.20 | k3.0 | r1.2 | 0.77105 | 0.082033 | 0.61 | 22.05 |
| b3 | p0.90 | t1.00 | k6.0 | r1.4 | 0.77065 | 0.061983 | 0.6 | 22.15 |
| b3 | p0.90 | t1.20 | k6.0 | r1.2 | 0.7704 | 0.072887 | 0.59 | 22.1 |
| b0 | p0.95 | t0.90 | k3.0 | r1.0 | 0.77035 | 0.075353 | 0.61 | 21.4 |
| b3 | p0.95 | t1.20 | k3.0 | r1.0 | 0.7699 | 0.088431 | 0.59 | 21.4 |
| b3 | p0.95 | t1.30 | k3.0 | r1.4 | 0.7697 | 0.075054 | 0.61 | 22.15 |
| b3 | p0.90 | t1.20 | k6.0 | r1.4 | 0.7696 | 0.063874 | 0.58 | 21.9 |
| b3 | p0.85 | t1.20 | k6.0 | r1.4 | 0.769 | 0.071613 | 0.58 | 22.2 |
| b3 | p0.90 | t1.00 | k6.0 | r1.0 | 0.7688 | 0.086077 | 0.6 | 22.25 |
| b3 | p0.85 | t1.20 | k3.0 | r1.4 | 0.7688 | 0.086384 | 0.6 | 22.35 |
| b3 | p0.90 | t1.10 | k3.0 | r1.4 | 0.76815 | 0.078478 | 0.6 | 22.2 |
| b3 | p0.85 | t1.20 | k3.0 | r1.2 | 0.76775 | 0.094988 | 0.6 | 22.5 |
| b3 | p0.95 | t1.30 | k6.0 | r1.4 | 0.76765 | 0.059208 | 0.6 | 22.1 |
| b3 | p0.85 | t1.00 | k3.0 | r1.2 | 0.76765 | 0.081014 | 0.6 | 21.65 |
| b3 | p0.85 | t1.20 | k6.0 | r1.0 | 0.767 | 0.092766 | 0.605 | 21.75 |
| b3 | p0.90 | t1.30 | k6.0 | r1.4 | 0.76665 | 0.079421 | 0.59 | 22.2 |
| b3 | p0.90 | t1.10 | k6.0 | r1.0 | 0.7666 | 0.090347 | 0.59 | 22.05 |
| b3 | p0.90 | t1.00 | k6.0 | r1.2 | 0.7666 | 0.069108 | 0.59 | 22.05 |
| b3 | p0.85 | t1.10 | k3.0 | r1.4 | 0.76635 | 0.082348 | 0.59 | 22.2 |
| b3 | p0.85 | t1.10 | k6.0 | r1.2 | 0.7661 | 0.078844 | 0.6 | 21.6 |
| b3 | p0.85 | t1.00 | k6.0 | r1.2 | 0.766 | 0.073958 | 0.59 | 22.6 |
| b3 | p0.85 | t1.10 | k6.0 | r1.0 | 0.7658 | 0.085669 | 0.59 | 21.85 |
| b0 | p0.95 | t0.80 | k3.0 | r1.0 | 0.76565 | 0.077898 | 0.6 | 21.5 |
| b0 | p0.95 | t0.80 | k3.0 | r1.2 | 0.7656 | 0.040014 | 0.6 | 20.9 |
| b3 | p0.85 | t1.30 | k6.0 | r1.4 | 0.76555 | 0.073249 | 0.6 | 22.1 |
| **b0** | **p0.85** | **t1.00** | **k3.0** | **r1.2** | **0.7655** | **0.036388** | **0.59** | **20.45** |
| b3 | p0.95 | t1.10 | k3.0 | r1.2 | 0.7653 | 0.091382 | 0.6 | 21.7 |
| b3 | p0.90 | t1.10 | k6.0 | r1.4 | 0.76515 | 0.068465 | 0.6 | 22.1 |
| b3 | p0.90 | t1.00 | k3.0 | r1.4 | 0.765 | 0.089751 | 0.6 | 21.6 |
| b0 | p0.90 | t1.10 | k3.0 | r1.0 | 0.7649 | 0.072978 | 0.6 | 21.45 |
| b3 | p0.85 | t1.20 | k6.0 | r1.2 | 0.76485 | 0.080018 | 0.6 | 22.45 |
| b3 | p0.90 | t1.20 | k3.0 | r1.2 | 0.7648 | 0.083571 | 0.6 | 22.6 |
| b3 | p0.85 | t1.00 | k3.0 | r1.4 | 0.7648 | 0.079301 | 0.59 | 22.15 |
| b3 | p0.85 | t1.00 | k6.0 | r1.4 | 0.7648 | 0.074135 | 0.59 | 22.2 |
| b3 | p0.90 | t1.30 | k6.0 | r1.0 | 0.76475 | 0.069082 | 0.58 | 22.1 |
| b0 | p0.85 | t1.00 | k3.0 | r1.0 | 0.76475 | 0.063778 | 0.6 | 21.4 |
| b0 | p0.85 | t1.00 | k6.0 | r1.0 | 0.7647 | 0.043986 | 0.6 | 21.05 |
| b3 | p0.90 | t1.00 | k3.0 | r1.2 | 0.7647 | 0.087533 | 0.59 | 22 |
| b3 | p0.90 | t1.10 | k3.0 | r1.0 | 0.7647 | 0.094374 | 0.58 | 21.75 |
| b3 | p0.85 | t1.00 | k3.0 | r1.0 | 0.7646 | 0.094856 | 0.59 | 21.7 |
| b3 | p0.90 | t1.30 | k3.0 | r1.4 | 0.76455 | 0.088426 | 0.6 | 22.55 |
| b0 | p0.95 | t1.20 | k3.0 | r1.2 | 0.76455 | 0.033463 | 0.61 | 20 |
| b0 | p0.85 | t1.20 | k3.0 | r1.0 | 0.76435 | 0.074973 | 0.6 | 21.25 |
| b3 | p0.95 | t1.30 | k6.0 | r1.2 | 0.76425 | 0.064431 | 0.6 | 22.25 |
| b3 | p0.95 | t1.00 | k3.0 | r1.4 | 0.76405 | 0.083046 | 0.59 | 22.4 |
| b3 | p0.95 | t1.00 | k6.0 | r1.2 | 0.76405 | 0.074841 | 0.6 | 22.2 |
| b3 | p0.85 | t1.30 | k3.0 | r1.4 | 0.76395 | 0.091574 | 0.59 | 22.7 |

| | | | | | | | | |
|---|---|---|---|---|---|---|---|---|
| b3 | p0.90 | t1.10 | k3.0 | r1.2 | 0.7639 | 0.081197 | 0.59 | 22.7 |
| b3 | p0.85 | t1.30 | k3.0 | r1.2 | 0.76375 | 0.088861 | 0.585 | 22.7 |
| b3 | p0.90 | t1.10 | k6.0 | r1.2 | 0.76375 | 0.075914 | 0.59 | 22.4 |
| b3 | p0.90 | t1.20 | k3.0 | r1.4 | 0.7636 | 0.071306 | 0.595 | 22.15 |
| b0 | p0.90 | t1.20 | k3.0 | r1.0 | 0.76345 | 0.06945 | 0.59 | 20.9 |
| b3 | p0.95 | t1.10 | k3.0 | r1.4 | 0.7633 | 0.081274 | 0.6 | 22.2 |
| b3 | p0.95 | t1.20 | k6.0 | r1.0 | 0.76325 | 0.088064 | 0.59 | 22.05 |
| b0 | p0.85 | t1.10 | k3.0 | r1.0 | 0.76325 | 0.058214 | 0.59 | 21 |
| b0 | p0.90 | t1.10 | k3.0 | r1.2 | 0.7627 | 0.037792 | 0.59 | 20.8 |
| b3 | p0.85 | t1.00 | k6.0 | r1.0 | 0.76265 | 0.091792 | 0.58 | 21.7 |
| b0 | p0.95 | t1.10 | k3.0 | r1.0 | 0.76265 | 0.072896 | 0.59 | 21.2 |
| b3 | p0.95 | t1.20 | k6.0 | r1.2 | 0.7625 | 0.076149 | 0.61 | 21.7 |
| b3 | p0.85 | t1.30 | k6.0 | r1.2 | 0.76245 | 0.067257 | 0.59 | 22 |
| b0 | p0.85 | t1.30 | k3.0 | r1.0 | 0.76225 | 0.056085 | 0.59 | 21.1 |
| b3 | p0.95 | t1.30 | k3.0 | r1.2 | 0.76195 | 0.082584 | 0.59 | 22.1 |
| b3 | p0.90 | t1.30 | k3.0 | r1.0 | 0.76145 | 0.090261 | 0.59 | 21.8 |
| b3 | p0.85 | t1.30 | k6.0 | r1.0 | 0.7614 | 0.083033 | 0.59 | 21.5 |
| b0 | p0.85 | t0.80 | k3.0 | r1.2 | 0.7613 | 0.040544 | 0.615 | 20.55 |
| greedy | | | | | 0.7612 | 0.098858 | 0.585 | 22.2 |
| b3 | p0.95 | t1.00 | k6.0 | r1.4 | 0.7611 | 0.073835 | 0.58 | 22 |
| b3 | p0.90 | t1.20 | k6.0 | r1.0 | 0.76105 | 0.07777 | 0.6 | 21.5 |
| b3 | p0.85 | t1.20 | k3.0 | r1.0 | 0.761 | 0.08298 | 0.6 | 22.05 |
| b0 | p0.90 | t0.90 | k3.0 | r1.2 | 0.761 | 0.041887 | 0.605 | 20.6 |
| b3 | p0.90 | t1.20 | k3.0 | r1.0 | 0.76095 | 0.093233 | 0.59 | 21.9 |
| b3 | p0.95 | t1.20 | k6.0 | r1.4 | 0.7609 | 0.067927 | 0.59 | 21.9 |
| b3 | p0.90 | t1.30 | k3.0 | r1.2 | 0.76065 | 0.089971 | 0.6 | 22.6 |
| b3 | p0.95 | t1.30 | k3.0 | r1.0 | 0.76065 | 0.089326 | 0.59 | 21.5 |
| b0 | p0.85 | t0.90 | k3.0 | r1.2 | 0.76045 | 0.043142 | 0.59 | 20.55 |
| b3 | p0.95 | t1.10 | k6.0 | r1.4 | 0.76035 | 0.076888 | 0.6 | 22.2 |
| b0 | p0.90 | t0.80 | k3.0 | r1.0 | 0.76015 | 0.076055 | 0.6 | 21.25 |
| b3 | p0.85 | t1.10 | k3.0 | r1.0 | 0.76 | 0.095941 | 0.59 | 22.2 |
| b0 | p0.90 | t1.00 | k3.0 | r1.0 | 0.7595 | 0.067128 | 0.6 | 21.55 |
| b3 | p0.95 | t1.10 | k6.0 | r1.2 | 0.7594 | 0.080542 | 0.59 | 22.2 |
| b0 | p0.85 | t0.90 | k3.0 | r1.0 | 0.7593 | 0.07734 | 0.6 | 21.2 |
| b0 | p0.90 | t0.90 | k6.0 | r1.0 | 0.75925 | 0.054728 | 0.6 | 21.1 |
| b0 | p0.90 | t0.90 | k3.0 | r1.0 | 0.7592 | 0.065855 | 0.59 | 20.9 |
| b3 | p0.95 | t1.10 | k6.0 | r1.0 | 0.7592 | 0.08592 | 0.59 | 22.2 |
| b3 | p0.95 | t1.30 | k6.0 | r1.0 | 0.7591 | 0.071843 | 0.58 | 21.85 |
| b3 | p0.95 | t1.10 | k3.0 | r1.0 | 0.75865 | 0.085117 | 0.58 | 21.5 |
| b3 | p0.95 | t1.20 | k3.0 | r1.4 | 0.7584 | 0.074848 | 0.595 | 22.5 |
| b0 | p0.85 | t0.80 | k3.0 | r1.0 | 0.75835 | 0.072269 | 0.6 | 21.8 |
| b3 | p0.85 | t1.10 | k6.0 | r1.4 | 0.75825 | 0.071193 | 0.595 | 22.05 |
| b0 | p0.90 | t0.80 | k3.0 | r1.2 | 0.7581 | 0.040474 | 0.6 | 20.65 |
| b0 | p0.90 | t1.20 | k3.0 | r1.2 | 0.7581 | 0.032783 | 0.605 | 20.35 |
| b0 | p0.85 | t1.20 | k3.0 | r1.4 | 0.758 | 0.024085 | 0.59 | 19.7 |
| b0 | p0.90 | t1.00 | k3.0 | r1.2 | 0.75795 | 0.042946 | 0.6 | 20.45 |
| b0 | p0.85 | t0.80 | k6.0 | r1.0 | 0.75785 | 0.063189 | 0.59 | 21.7 |
| b0 | p0.90 | t0.80 | k6.0 | r1.0 | 0.7578 | 0.056882 | 0.59 | 21.2 |
| b0 | p0.95 | t0.90 | k3.0 | r1.2 | 0.75765 | 0.037636 | 0.605 | 20.3 |
| b0 | p0.85 | t1.20 | k3.0 | r1.2 | 0.7576 | 0.045269 | 0.6 | 20.45 |
| b3 | p0.95 | t1.00 | k3.0 | r1.0 | 0.757 | 0.098715 | 0.59 | 22.2 |
| b0 | p0.90 | t1.30 | k3.0 | r1.0 | 0.7566 | 0.062032 | 0.6 | 21 |
| b0 | p0.95 | t1.30 | k3.0 | r1.0 | 0.75655 | 0.068489 | 0.605 | 20.5 |
| b0 | p0.95 | t1.10 | k3.0 | r1.2 | 0.75625 | 0.031058 | 0.59 | 20.2 |
| b3 | p0.90 | t1.00 | k3.0 | r1.0 | 0.75615 | 0.092407 | 0.59 | 22.2 |

| | | | | | | | | |
|---|---|---|---|---|---|---|---|---|
| b3 | p0.90 | t1.30 | k6.0 | r1.2 | 0.7556 | 0.073403 | 0.59 | 22.05 |
| b0 | p0.85 | t1.10 | k3.0 | r1.2 | 0.75555 | 0.033202 | 0.6 | 20.5 |
| b0 | p0.90 | t0.90 | k6.0 | r1.2 | 0.75515 | 0.016035 | 0.6 | 20.4 |
| b0 | p0.95 | t0.80 | k3.0 | r1.4 | 0.75515 | 0.032418 | 0.59 | 20 |
| b0 | p0.85 | t1.30 | k3.0 | r1.2 | 0.75495 | 0.032353 | 0.595 | 19.6 |
| b0 | p0.85 | t0.80 | k6.0 | r1.2 | 0.75495 | 0.027836 | 0.6 | 20.75 |
| b0 | p0.85 | t1.30 | k6.0 | r1.0 | 0.7542 | 0.046825 | 0.585 | 19.9 |
| b0 | p0.95 | t1.00 | k3.0 | r1.2 | 0.7542 | 0.040389 | 0.6 | 20.45 |
| b0 | p0.95 | t1.00 | k3.0 | r1.0 | 0.75365 | 0.070251 | 0.59 | 20.7 |
| b3 | p0.95 | t1.00 | k6.0 | r1.0 | 0.75345 | 0.079371 | 0.59 | 21.8 |
| b0 | p0.95 | t0.80 | k6.0 | r1.0 | 0.7534 | 0.052797 | 0.6 | 20.8 |
| b0 | p0.85 | t0.90 | k6.0 | r1.2 | 0.7532 | 0.027534 | 0.59 | 20.05 |
| b0 | p0.85 | t1.20 | k6.0 | r1.0 | 0.7527 | 0.035154 | 0.595 | 20.05 |
| b0 | p0.95 | t0.90 | k6.0 | r1.0 | 0.7523 | 0.052248 | 0.6 | 20.15 |
| b0 | p0.85 | t0.80 | k3.0 | r1.4 | 0.7519 | 0.032417 | 0.59 | 19.8 |
| b0 | p0.85 | t1.00 | k6.0 | r1.2 | 0.7518 | 0.019711 | 0.6 | 20.3 |
| b0 | p0.90 | t1.10 | k3.0 | r1.4 | 0.75145 | 0.030054 | 0.6 | 19.85 |
| b0 | p0.95 | t1.30 | k3.0 | r1.2 | 0.75145 | 0.028702 | 0.6 | 20 |
| b0 | p0.90 | t1.00 | k6.0 | r1.2 | 0.75085 | 0.021128 | 0.6 | 19.2 |
| b0 | p0.95 | t0.80 | k6.0 | r1.2 | 0.75085 | 0.020436 | 0.58 | 20 |
| b0 | p0.85 | t1.10 | k6.0 | r1.2 | 0.7506 | 0.01807 | 0.6 | 19.55 |
| b0 | p0.90 | t0.80 | k6.0 | r1.2 | 0.7503 | 0.023444 | 0.6 | 20 |
| b0 | p0.95 | t0.90 | k3.0 | r1.4 | 0.75 | 0.033901 | 0.59 | 19.65 |
| b0 | p0.85 | t0.90 | k6.0 | r1.0 | 0.74985 | 0.055786 | 0.59 | 21.7 |
| b0 | p0.90 | t1.10 | k6.0 | r1.0 | 0.7494 | 0.040637 | 0.58 | 20 |
| b0 | p0.85 | t0.90 | k3.0 | r1.4 | 0.7494 | 0.038356 | 0.6 | 20.15 |
| b0 | p0.90 | t1.10 | k6.0 | r1.2 | 0.74925 | 0.014442 | 0.59 | 19.6 |
| b0 | p0.90 | t1.00 | k3.0 | r1.4 | 0.7491 | 0.029877 | 0.6 | 19.5 |
| b0 | p0.90 | t0.90 | k3.0 | r1.4 | 0.7491 | 0.029974 | 0.59 | 19.8 |
| b0 | p0.90 | t1.20 | k3.0 | r1.4 | 0.74905 | 0.025708 | 0.6 | 19 |
| b0 | p0.90 | t1.30 | k3.0 | r1.2 | 0.74905 | 0.037212 | 0.595 | 20 |
| b0 | p0.95 | t1.30 | k6.0 | r1.0 | 0.74855 | 0.029211 | 0.58 | 19.35 |
| b0 | p0.95 | t1.20 | k3.0 | r1.0 | 0.7484 | 0.064294 | 0.59 | 20.5 |
| b0 | p0.85 | t1.00 | k3.0 | r1.4 | 0.7481 | 0.030258 | 0.61 | 19.5 |
| b0 | p0.90 | t1.30 | k6.0 | r1.0 | 0.7479 | 0.039046 | 0.58 | 19.8 |
| b0 | p0.85 | t1.20 | k6.0 | r1.2 | 0.7473 | 0.018512 | 0.59 | 19.3 |
| b0 | p0.85 | t1.10 | k3.0 | r1.4 | 0.7467 | 0.029222 | 0.595 | 19 |
| b0 | p0.95 | t0.90 | k6.0 | r1.2 | 0.74645 | 0.018903 | 0.58 | 19 |
| b0 | p0.90 | t0.80 | k3.0 | r1.4 | 0.7459 | 0.034532 | 0.58 | 20.05 |
| b0 | p0.90 | t1.00 | k6.0 | r1.0 | 0.7459 | 0.04542 | 0.58 | 20.45 |
| b0 | p0.90 | t1.30 | k3.0 | r1.4 | 0.74585 | 0.025511 | 0.595 | 19.05 |
| b0 | p0.85 | t0.90 | k6.0 | r1.4 | 0.74505 | 0.018003 | 0.58 | 19.05 |
| b0 | p0.85 | t1.10 | k6.0 | r1.0 | 0.74495 | 0.047379 | 0.59 | 20.2 |
| b0 | p0.95 | t1.10 | k3.0 | r1.4 | 0.7448 | 0.028739 | 0.585 | 19.4 |
| b0 | p0.90 | t1.20 | k6.0 | r1.0 | 0.7446 | 0.039695 | 0.58 | 20 |
| b0 | p0.95 | t1.10 | k6.0 | r1.0 | 0.74435 | 0.038183 | 0.58 | 19.55 |
| b0 | p0.85 | t1.30 | k3.0 | r1.4 | 0.74405 | 0.028199 | 0.585 | 19.7 |
| b0 | p0.95 | t1.00 | k3.0 | r1.4 | 0.74385 | 0.025351 | 0.59 | 19.1 |
| b0 | p0.95 | t1.30 | k3.0 | r1.4 | 0.7432 | 0.028738 | 0.58 | 19.6 |
| b0 | p0.95 | t1.20 | k3.0 | r1.4 | 0.7429 | 0.029231 | 0.6 | 19.1 |
| b0 | p0.95 | t1.00 | k6.0 | r1.0 | 0.74215 | 0.033477 | 0.58 | 19.7 |
| b0 | p0.85 | t1.00 | k6.0 | r1.4 | 0.7408 | 0.013787 | 0.57 | 18.7 |
| b0 | p0.85 | t1.30 | k6.0 | r1.2 | 0.73985 | 0.016491 | 0.57 | 19 |
| b0 | p0.90 | t0.90 | k6.0 | r1.4 | 0.7394 | 0.013402 | 0.58 | 19.3 |
| b0 | p0.95 | t1.20 | k6.0 | r1.2 | 0.7393 | 0.011679 | 0.59 | 18.05 |

| | | | | | | | | |
|---|---|---|---|---|---|---|---|---|
| b0 | p0.90 | t1.30 | k6.0 | r1.2 | 0.739 | 0.013721 | 0.58 | 18.2 |
| b0 | p0.85 | t0.80 | k6.0 | r1.4 | 0.7389 | 0.019544 | 0.59 | 19.45 |
| b0 | p0.95 | t1.00 | k6.0 | r1.2 | 0.73885 | 0.015533 | 0.585 | 18.8 |
| b0 | p0.90 | t0.80 | k6.0 | r1.4 | 0.73845 | 0.017694 | 0.59 | 19.45 |
| b0 | p0.95 | t1.10 | k6.0 | r1.2 | 0.7369 | 0.01298 | 0.57 | 18.6 |
| b0 | p0.95 | t0.80 | k6.0 | r1.4 | 0.73545 | 0.011923 | 0.58 | 18.45 |
| b0 | p0.95 | t1.20 | k6.0 | r1.0 | 0.73495 | 0.035326 | 0.58 | 19.6 |
| b0 | p0.85 | t1.10 | k6.0 | r1.4 | 0.7337 | 0.014377 | 0.57 | 18.35 |
| b0 | p0.90 | t1.20 | k6.0 | r1.2 | 0.72965 | 0.015058 | 0.59 | 19.5 |
| b0 | p0.85 | t1.20 | k6.0 | r1.4 | 0.72595 | 0.014126 | 0.565 | 18.45 |
| b0 | p0.95 | t0.90 | k6.0 | r1.4 | 0.72515 | 0.013653 | 0.57 | 18.5 |
| b0 | p0.90 | t1.10 | k6.0 | r1.4 | 0.72495 | 0.010673 | 0.57 | 17.9 |
| b0 | p0.95 | t1.30 | k6.0 | r1.2 | 0.72445 | 0.012223 | 0.57 | 18.15 |
| b0 | p0.95 | t1.00 | k6.0 | r1.4 | 0.7242 | 0.010586 | 0.57 | 18.35 |
| b0 | p0.85 | t1.30 | k6.0 | r1.4 | 0.72215 | 0.012042 | 0.57 | 17.8 |
| b0 | p0.90 | t1.00 | k6.0 | r1.4 | 0.72095 | 0.016106 | 0.57 | 18.4 |
| b0 | p0.95 | t1.10 | k6.0 | r1.4 | 0.7174 | 0.008668 | 0.57 | 17.9 |
| b0 | p0.90 | t1.20 | k6.0 | r1.4 | 0.7161 | 0.011932 | 0.56 | 17.7 |
| b0 | p0.95 | t1.20 | k6.0 | r1.4 | 0.7144 | 0.01096 | 0.57 | 17.4 |
| b0 | p0.90 | t1.30 | k6.0 | r1.4 | 0.71295 | 0.010127 | 0.57 | 17.4 |
| b0 | p0.95 | t1.30 | k6.0 | r1.4 | 0.7016 | 0.011304 | 0.55 | 16.9 |

* Comparison of various generation configurations tested for translating from 3Di sequences to amino acid sequences for proteins in the validation set (3Di→AA). Metrics were computed between *groundtruth* (3D structure derived from AlphaFold2 predictions[16]) and *predictions* (ESMFold[20] predictions of ProstT5-generated sequences). The columns show a) the number of beams (b0=no beam, b3=3 beams), b) the p-value (p0.9 means p=0.9), c) the temperature (t0.80 refers to temperature=0.8), d) the k-value (k3.0 means k-value=3), e) the repetition penalty (r1.2 means repetition-penalty=1.2), f), g), h) and i) show the medians of lDDT[15], entropy (KL-divergence between AA distribution in UniProt[21] and generated sequences), TM-score[22], and sequence similarity, respectively. For comparison, we also added the performance of simple greedy sampling (always picking the next token based on highest probability). For the ProstT5 paper we chose the configuration highlighted in bold.
